# TubULAR: Tracking *in toto* deformations of dynamic tissues via constrained maps

**DOI:** 10.1101/2022.04.19.488840

**Authors:** Noah P. Mitchell, Dillon J. Cislo

## Abstract

A common motif in biology is the arrangement of cells into tube-like sheets, which further transform into more complex shapes. Traditionally, analysis of the dynamic surfaces of growing tissues has relied on inspecting static snapshots, live imaging of cross-sections, or tracking isolated cells in 3D. However, capturing the interplay between in-plane and out-of-plane behaviors requires following the full surface as it deforms and integrating cell-scale motions into collective, tissue-scale deformations. The advent of methods for whole-organ live imaging ushers the need for *in toto* analysis methods to understand these collective tissue dynamics. Here, we introduce an approach that builds *in toto* maps of surface deformation by following tissue parcels in the material frame of reference. Mapping the full 3D surface to 2D images such that the tissue motion in 2D is minimal enables the user to easily follow the tissue and discern its 3D motion. We then provide a computational framework for linking in-plane and out-of-plane behaviors and decomposing complex deformation maps into elementary contributions. The Tube-like sUrface Lagrangian Analysis Resource (TubULAR) provides an open-source MATLAB implementation whose functionality is accessible either as a standalone toolkit or as an extension of the ImSAnE package used in the developmental biology community. We underscore the power of our approach by analyzing shape change in the embryonic Drosophila midgut and beating zebrafish heart. Following deformations in the tissue/material frame reveals the signatures of tissue flow and a reduced-dimensional mode decomposition of the dynamics. The method naturally generalizes to *in vitro* and synthetic systems and provides ready access to the mechanical mechanisms relating genetic patterning to organ shape change.

## INTRODUCTION

In the morphogenesis of thin tissues, the interplay between mechanical forces, cellular fates, and physiological function determines dynamic patterns of shape change. In epithelia [1, 2], visceral organs [3, 4], vasculature [5], elastic shells [6], and whole organisms alike [7], tube-like surfaces deform in 3D space, contracting and dilating inplane while bending out-of-plane in a coupled fashion. Understanding the dynamic mechanisms by which the systems change shape requires not only capturing instantaneous motion in 3D, but also following the material as it deforms to build so-called ‘Lagrangian’ measures of tissue deformation [8, 9]. Furthermore, decomposing that motion into physically meaningful components enables insights into the processes generating organ shape and function [4, 10–12].

Much of of our understanding of morphogenesis built during the last century has relied on qualitative analysis of 2D live imaging and static snapshots of dynamic growth processes in 3D [13–15]. Emerging computational approaches in the life sciences have enabled quantitative characterizations that often challenge traditional assumptions and clarify the complex relationship between gene expression patterns, physical forces, and tissue geometry [9, 16–22]. In a particularly fruitful methodological advance, the community has applied ‘tissue cartography’ methods that map curved tissues to a planar representation, dramatically reducing the computational power required to store and analyze 3D data [23–26]. This advance has facilitated insights in a wide variety of systems including fly wings [27], eyes [28], egg chambers [29], ascidian vasculature [30], zebrafish endoderm [23], and mouse intestines [31].

While these methods are sufficient to track tissue motion within static geometries or in local patches [24], *in toto* measurements of tissue deformation in complex, dynamic geometries have remained elusive. Here we propose automated methods for registering dynamic tube-like surfaces across time – with arbitrarily complex geometry – and classifying the signatures of tissue deformation underlying organ-scale shape change. As shown in Fig. 1, this provides a framework for automatically tracing the dynamics of complex shapes and facilitates cell tracking on contorting 3D tissue surfaces. This framework then naturally decomposes tissue-frame measurements for interpretation, handling all computational subtleties that arise from the surface’s curvature and bending.

**FIG. 1.**
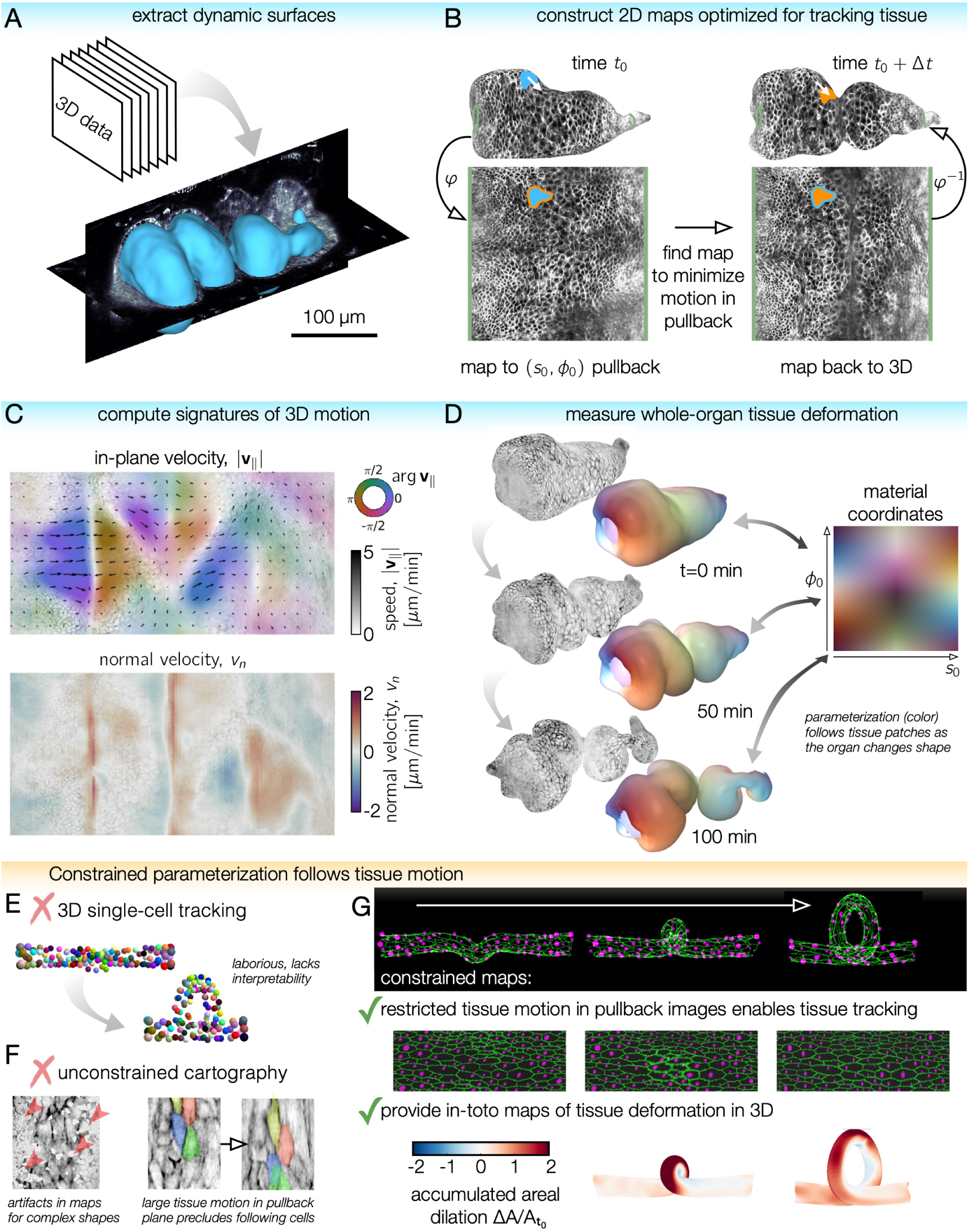
TubULAR is a toolkit for following the dynamics of evolving tube-like surfaces, such as visceral organs and *in vitro* biological surfaces. *(A)* TubULAR first extracts dynamic surfaces of interest from volumetric datasets. *(B)* Whole-surface parameterization using a generalized cylindrical coordinate system facilitates tracking tissue deformations. Since pullback images of adjacent timepoints are optimized to be nearly stationary in the parameterization space, 3D tissue velocities (white arrow) are obtained by linking the pre-image of the pullback coordinates across timepoints. *(C)* By extracting in-plane and out-of-plane tissue motion, TubULAR decomposes the underlying motion, here shown by a 2D pullback representation of the in-plane tissue velocity **v**_∥_ (colored quiverplot) and the normal motion, *v*_*n*_ (red for inward velocity, blue for outward). *(D)* Finally, the toolkit tracks tissue deformations over time in the tissue’s frame of reference (the Lagrangian frame). Here, a whole organ is colored by the location of each tissue parcel in its intrinsic material coordinate system (*s*_0_, *ϕ*_0_). Patches retain their original color as they move, stretch, and bend. *(E)* Tracking individual cells typically involves laborious manual input and requires interpretation for translating into tissue-scale deformation patterns. Cell identities are colored from an *in silico* example of cells on a coiling tube. *(F)* Cartographic projections using previously-published ImSAnE methods fail for complex and dynamic shapes such as the folding midgut. (Left) Red arrows mark several points where the multivalued surface returns image artifacts using ImSAnE’s cylinderMeshWrapper. (Right) Motion of the mapped cells in the pullback plane is large for adjacent timepoints without placing additional constraints on the pullback map. *(G)* An *in silico* dataset maps tissue to a series of images which change little over time. Tracking the motion in 2D with ease then allows measurement of the tissue deformation across the full tube in 3D, here shown for the accumulated dilatational strain.

By applying this approach to the developing digestive tract in the embryonic fly *Drosophila melanogaster* [4, 32–34] and the beating heart of the embryonic zebrafish *Danio rerio* [3, 12, 35–37], we extract the full deformation fields and relate signatures of in-plane and out-of-plane tissue deformation. Despite the complexity of their motion, we obtain simple geometric descriptions underlying shape change.

## IMPLEMENTATION AND APPLICATION TO A MODEL ORGAN

Contemporary microscopy methods generate volumetric data, wherein each voxel carries a (potentially multichannel) intensity measured at a specific location in the sample [38]. At the same time, many biological processes harness quasi-2D, thin tissues or interfaces to sculpt complex 3D forms. To probe the interplay between in-plane interactions and out-of-plane dynamics in such systems, we must extract the tissue surface, track motion within the surface as it deforms, and decompose the resulting motion into signatures of deformation.

We package this functionality in the Tube-like sUrface Lagrangian Analysis Resource (TubULAR). TubULAR is publically available on GitHub, with documentation and example scripts at https://npmitchell.github.io/tubular/. The package includes (and uses) independent toolkits for surface visualization (TexturePatch), conformal mapping (RicciFlow), and discrete exterior calculus (DECLab). A typical workflow passes through (1) level sets segmentation for surface detection, (2) generation of a pullback representation for surface parameterization and visualization, (3) flow analysis using TubULAR’s discrete exterior calculus implementation, and, if appropriate, (4) mode decomposition of the dynamics. In addition to the standalone toolkit, we have incorporated the core functionality of TubULAR within the ImSAnE environment [24], with updates available at https://github.com/npmitchell/imsane.

### From volumetric data to dynamic textured surfaces

We first set out to identify and extract 2D surfaces of interest from 3D data. For the tube-like shape of the systems we address here, TubULAR segments the data space into an ‘inside’ (i.e. everything within the tube) and an ‘outside’ (i.e. everything outside the tube), modulo the potentially necessary inclusion of virtual ‘end caps’ to close off the interior of an open tube [39]. As illustrated in Fig. 1A, level set methods are a robust and powerful tool used to dynamically segment data volumes into complex watertight geometries whose boundaries are defined as the null contours of a signed distance function *u*(**x**). The problem of surface detection is thereby reduced to optimizing the signed distance function so that *u*(**x**) = 0 matches the tissue surface.

Once a level set partitions the data volume for a given time point, we the point cloud defined by the voxels on the boundary of the segmented region to generate a smooth mesh triangulation. The result is a dynamic set of surfaces tracing the tube-like surface over time, as illustrated in Fig. 1B-D. Users may alternately generate surfaces via other software (such as Imaris), then use TubULAR for subsequent analysis. A toolkit for rendering the data onto the surface (TexturePatch) is included and implemented within TubULAR – and is also functional as a standalone tool (Fig. 1B-D).

### Constrained surface parameterization enables tracking surface dynamics in the material frame

Understanding the ways in which shape dynamics couple to biological processes such as cell shape change, intercalation, intracellular patterning, and gene expression requires the ability to identify and follow patches of tissue as they move and deform. On its own, the previous surface extraction step provides an instantaneous description of the surface geometry, but does not identify how particular patches of cells move and deform from time point to time point. Generating a consistent, time-dependent coordinate system taking into account tissue flow and deformation presents a considerable technical challenge, particularly in the presence of dramatic shape change. In the language of continuum mechanics, we must advance from an *Eulerian* description, wherein dynamical observables are measured at particular locations in space, to a *Lagrangian* description, wherein the dynamics – whether of deformation, anisotropy, morphogen concentration, or any another observable – are queried by following material parcels along the surface as morphogenesis proceeds [40].

As illustrated in Fig. 1B and Fig. 2, we build a parameterization scheme such that the motion of the cells (or other objects) in the pullback representation move as little as possible. This enables us to easily follow a virtual representation of in-plane deformations in pullback space, which we can then project into 3D to follow the true motion. To this end, TubULAR first cartographically maps the surface to the plane – defining a material frame of reference – then stabilizes virtual motion of the material in the pullback plane. The result is a dynamic map *φ*(*t*) mapping the dynamic surface to a fixed material frame of reference, with minimal movement of the tissue over time in the pullback images (Fig. 2).

**FIG. 2.**
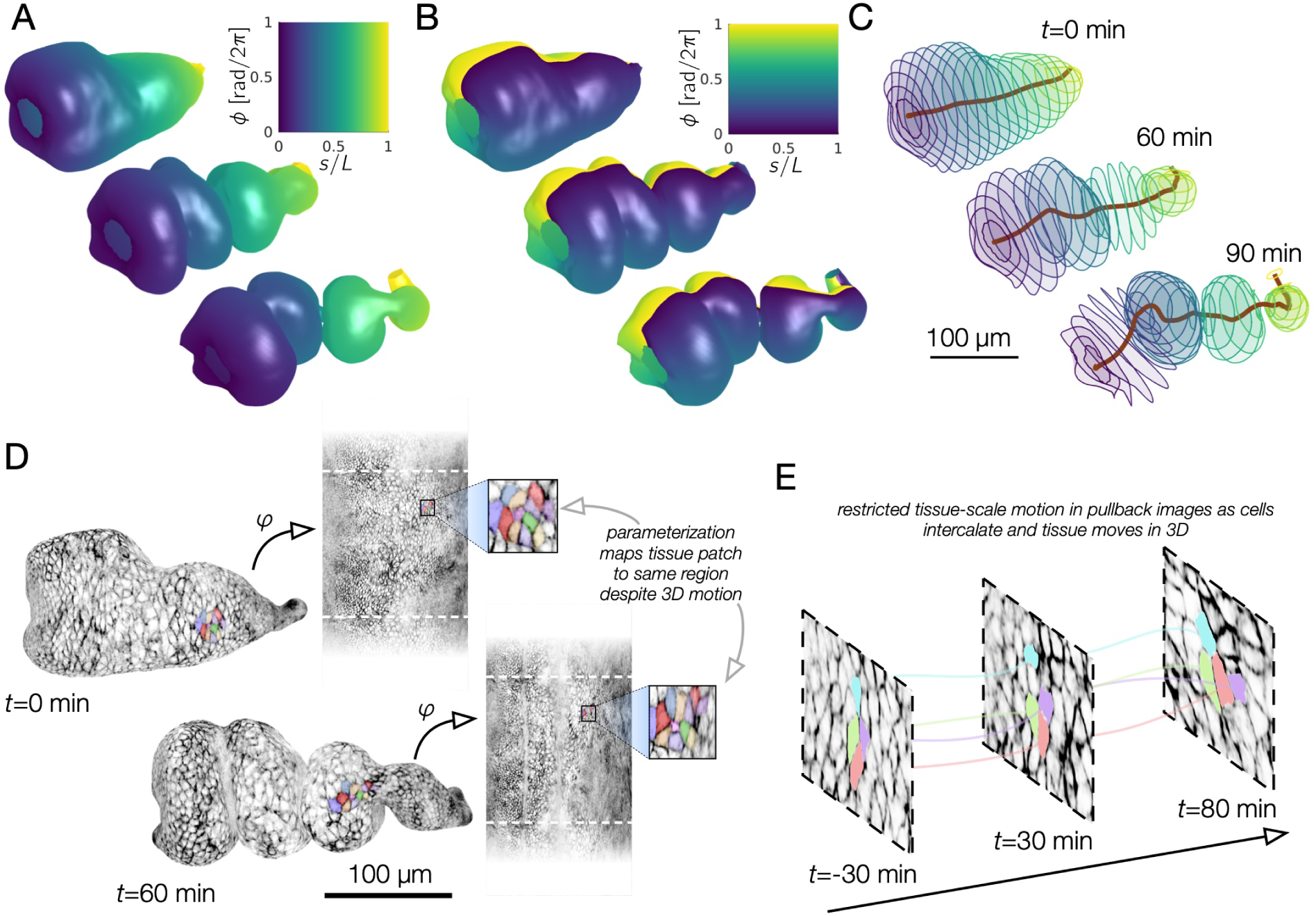
Coordinate parameterization follows 3D deformations of the evolving surface, enabling quantification of geometric dynamics, shown here for the fly midgut as it folds into compartments. *(A)* Constructing a dynamic map to fixed material coordinates, labels the midgut’s intrinsic longitudinal axis, even as the surface deforms into a convoluted shape. A 2D pullback representation of the material frame parameterizes the longitudinal axis by color as indicated. *(B)* Dynamic mapping to fixed material coordinates tracks circumferential motion of the tissue. *(C)* Computational sectioning of the parameterized surface shows circumferential disks of the organ sampled evenly along its length. This construction provides a natural, system-spanning centerline and a measure of effective radius along the surface. *(D)* Despite large tissue movement in 3D, tissue motion is restricted in the pullback images. *(E)* While tissue motion is small, individual cells may move and exchange neighbors. Measuring the difference between tissue strain rate and intercalation rate gives insights into the morphogenetic process (see Supplementary Information).

We first map the mesh at a single reference timepoint into the plane to define the material frame. To do so, we cut small ‘endcaps’ at the poles and create a virtual seam along the long axis so that the now-cylindrical mesh topology can be ‘unrolled’ into the plane. To facilitate the analysis of anisotropic tissue deformations later in the pipeline, this mapping into the plane is *conformal* – i.e. a totally isotropic, angle-preserving map. This conformal map is constructed by discrete Ricci flow [41] (included as a custom package within TubULAR, also a fully independent, standalone package) or can be approximated quickly via minimizing the Dirichlet ‘energy’ of the mapping [42] (see also Supplementary Information).

Finally, we advance from this initial map to follow the tissue motion. We do so by adjusting the mapping to the plane at other timepoints *φ*(*t*) so that any tissue patches are immobilized in the pullback image. As detailed in the Supplementary Information, we found that a four-step approach to constructing this map provides a balance between stringent motion minimization and stability across many timepoints: *φ*(*t*) = *J* ◦Φ◦*Z* ◦*f*, where *f* is a conformal map, *Z* adjusts the longitudinal axis, Φ minimizes motion along the circumferential axis, and *J* accounts for residual motion through optical correlation of adjacent pullback images. We are then able to follow extreme deformations of the tissue in 3D space simply by reading off the dynamic inverse map *φ*^−1^(*t*). Fig. 2A-C shows this parameterization scheme applied to the Drosophila midgut, with tracked surfaces colored by the longitudinal (Fig. 2A) and circumferential (Fig. 2B) coordinates of the tissue as it deforms in space. The resulting parameterization can be conceptualized as tracking the motion and rotations of deforming polygons arranged along a deforming, system-spanning centerline (Fig. 2C).

Fig. 2D-E shows that tracked cells in the corresponding pullback images move little despite large deformations of the organ surface in the Drosophila midgut. During embryonic development, the midgut closes into a tube composed of a monolayer of endoderm surrounded by a thin net of muscle cells [34, 43–46]. Subsequently, the midgut forms a constriction midway along its length, then two more constrictions subdividing the organ into four chambers [4, 32]. We apply our method to this surface and track cells in the pullback plane. Since the method rectifies tissue-scale motion, but not necessarily individual cells’ motion, we observe cell intercalation events such as those shown in Fig. 2E. Separating out the effects of tissue motion and cell intercalations has given insights into multiple mechanisms of morphogenesis in planar tissues [18, 47]. Here, this follows naturally from our approach in organs with complex shapes. The Supplementary Information details an example of this decomposition.

### Covariant measures of motion access signatures of deformation

By constructing material pathlines, we have already obtained velocity vectors of the tissue defined over the surface and over time. In order to *interpret* these tissue flows, we now decompose the velocity fields into their underlying components. A given parcel can move both along the surface and normal to the surface, and separating these motions is needed to parse whether tissue is rotating or shearing, contracting or dilating, and protruding or ingressing – and to find the spatiotemporal pattern in which these motifs occur. After computing a Lagrangian reference frame for the tissue, the default implementation in a TubULAR pipeline therefore (1) extracts the in-plane divergence and the local rotation rate of the tissue velocity, (2) relates the in-plane divergence to the out-of-plane motion *v*_*n*_ to determine the rate of areal growth across the surface, and (3) measure area-preserving shear deformations in the tissue.

Dealing with velocity fields on curved surfaces requires certain computational care: parallel lines cross and diverge, and the orientation of a cell may change by simply traveling along ‘straight’ lines (geodesics). In TubULAR, our calculations therefore rest on an implementation of the discrete exterior calculus (DEC) formalism [48, 49]. Signals are represented as discrete differential forms on the simplicial structure of the triangulated mesh which yield natural definitions for linear differential operators. These basic operations can be combined together to form covariant representations of more complicated differential operators, including the divergence, curl, and Laplacian operators that are a part of TubULAR’s default workflow. These operators can be directly applied to geometric data (e.g. surface curvature), kinematic data (e.g. surface velocities), and beyond (e.g. surface data intensity, surface data anisotropy fields etc.) to understand the ways in which spatiotemporal variation in observable fields (such as gene expression or myosin anisotropy) generate 3D shape change.

In order to make these methods accessible to the broadest possible audience, we have packaged all of this functionality within DECLab, a simple and flexible framework for discrete geometry processing. It is included with TubULAR and also functions as a standalone toolkit. No deep knowledge of differential geometry or exterior calculus is necessary to use our implementation (see Supplementary Information and online documentation).

Fig. 3 displays examples of these calculations applied to the developing Drosophila midgut. Whole-organ measurements of the tangential velocity are represented in the 2D pullback coordinates for snapshots of a representative embryo in Fig. 3A, with normal velocities shown in Fig. 3B. Further processing via DEC of the in-plane velocity fields shows localized sinks in the flow (∇ · **v**_∥_ *<* 0) near constrictions, as shown in Fig. 3C.

**FIG. 3.**
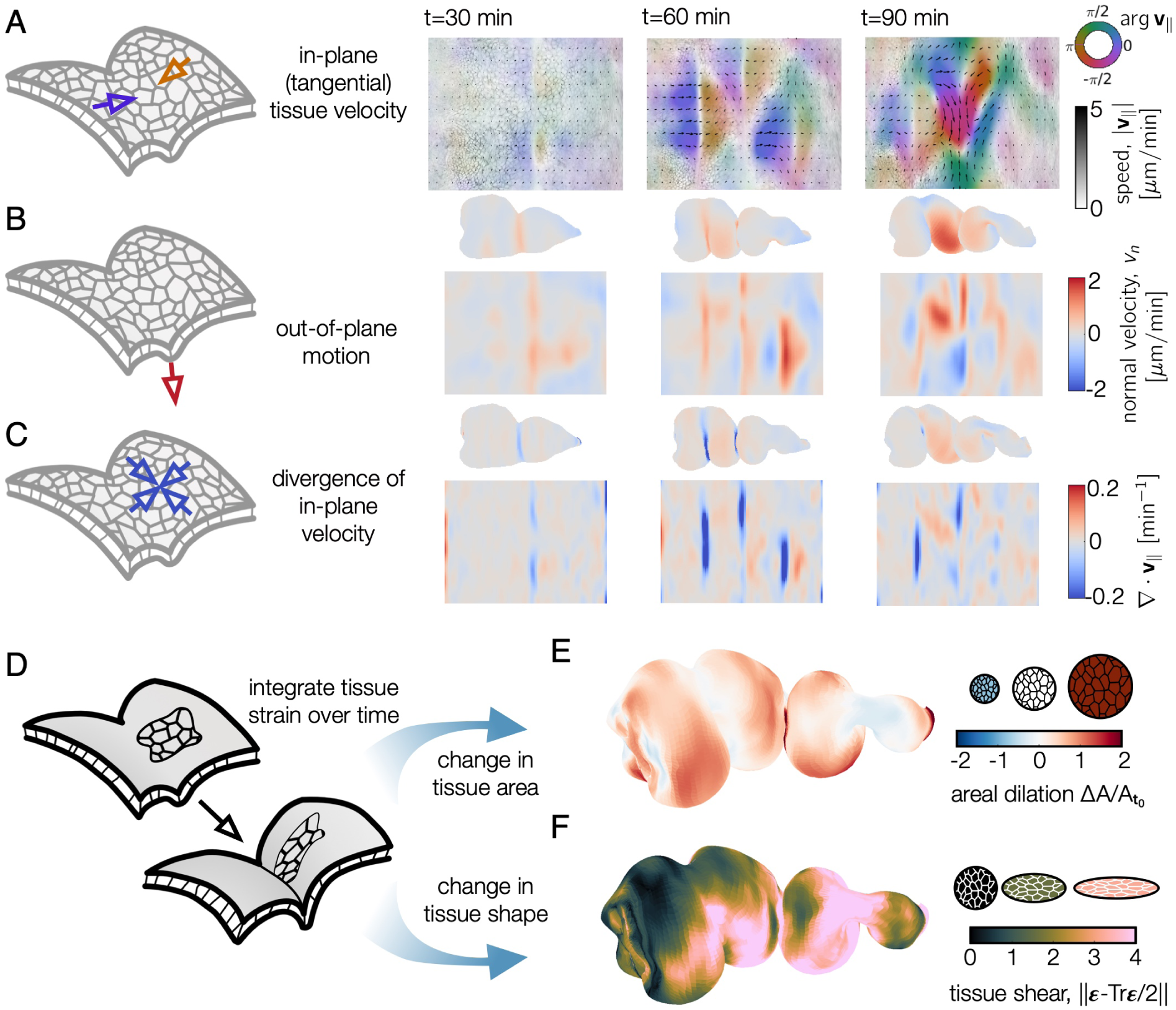
Dynamic maps to the material frame yield covariant measures of tissue velocity and deformation in 3D. Tissue velocities are decomposed into in-plane (tangential) and out-of-plane (normal) components. *(A)* 2D pullback representation of in-plane (tangential) tissue motion shows a line defect near the middle constriction, then defects at the anterior and posterior constrictions, followed by increasingly complex in-plane flows. The orientation of flow relative to the longitudinal axis *ŝ* is denoted by color and the magnitude of the motion reflected by opacity and arrow length. *(B)* The underlying out-of-plane velocity *v*_*n*_ is positive (inwards) near constrictions. *(C)* DEC computation of the divergence of the in-plane velocity ∇ · **v**_∥_ shows patterns of sinks in the constrictions and sources in the chambers’ lobes, in synchrony with the out-of-plane deformation. *(D-F)* Following material pathlines in 3D through time returns measures of integrated strain. These decompose into dilatational (area-changing) and deviatoric (shape-changing) components. *(E)* The dilatational strain shows areal growth in the chambers but less area change near constrictions for a midgut 90 minutes after the onset of the first constriction. *(F)* The deviatoric strain shows strong tissue shear near each constriction at the same timepoint.

### Lagrangian measures of time-integrated tissue strain

Endowing the evolving surface with a set of Lagrangian coordinates induces the construction of a *material metric*. The metric tensor, **g**(*t*), is a geometric object enabling the measurement of distances and angles between points on the surface. Unlike in flat geometries, here lengths and angles deform under the surface motion not only due to gradients in the tangential velocity, but also due to normal motion in curved regions of the tissue (see Supplementary Information). These changes are captured by the *rate-of-deformation* tensor, which is simply the time rate of change of the material metric, *d***g**(*t*)*/dt*, and can be constructed directly from the tissue velocity fields and surface curvatures. We can then integrate the rate-of-deformation tensor along pathlines to construct a Lagrangian measurement of cumulative tissue strain:

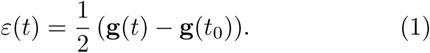

As illustrated in Fig. 3D-F, the result is decomposed into both an isotropic area change and an anisotropic shear deformation. The midgut shows increase in tissue area in the lobe of each chamber and only a slight decrease in tissue area near each constriction (Fig. 3E). These area changes result from accumulation of the small but persistent areal strain rate pattern (see Supplementary Information). The anisotropic shape-changing deformation indicative of convergent extension, in contrast, is strongest near each constriction (Fig. 3F). These integrated strain measures enable calculation of the rate of oriented cell intercalations, as detailed in the Supplementary Information.

### Field decomposition simplifies complicated deformations

Frequently, seemingly complicated patterns of motion can be decomposed into a sum of contributions from simpler components. This strategy has been successfully adopted in various contexts within morphogenesis, including zebrafish gastrulation [50] and vertebrate limb generation [51]. The TubULAR package deconstructs complex observable fields into elementary components at multiple stages of analysis.

We include the capability to decompose surface vector fields (e.g. material tissue velocities) into their geometric components. Our implementation generates a Helmholtz-Hodge decomposition of tangential vector fields [49], i.e. a decomposition into dilatational, rotational, and harmonic parts. This procedure determines the relative contribution of the geometric signatures of the in-plane flow to the out-of-plane deformation.

TubULAR also constructs mode decompositions of arbitrary tensor fields on the surface. This functionality comes in two forms. First, our DEC implementation decomposes signals onto a basis of the eigenfunctions of the discrete Laplace-Beltrami operator [52]. More simply, this compares the relative importance of longwavelength modes (with smooth spatial variation) and short-wavelength modes (with rapid spatial variation) in generating the observed signals. Second, we include functions to decompose signals using principal component analysis (PCA) [53], as demonstrated in the next section. This type of analysis lets users extract more general patterns of motion that contribute most strongly to the variance observed across time or across datasets.

## DECOMPOSING DEFORMATIONS DURING HEART MORPHOGENESIS

Pumping circulatory organs such as the mammalian heart are nearly omnipresent in animals. To demonstrate the generality of our method to cylic organ deformations, we analyzed a beating zebrafish heart during during the second day of development post fertilization. *In toto* imaging of the heart relied on light-sheet illumination of a transgenic Tg(*cmlc2:eGFP*) embryo expressing GFP in cardiomyocytes [54]. These data were taken using a temporal supperresolution approach, in which acquisition was synchronized with the beating of the developing heart to build volumetric data at 11 equally-spaced phases of the heart beat cycle [37]. The 3D shape of the beating heart is shown for a set of illustrative time points in Fig. 4A. Passing the volumetric data through TubULAR returns covariant measures of in-plane and out-of-plane deformation.

**FIG. 4.**
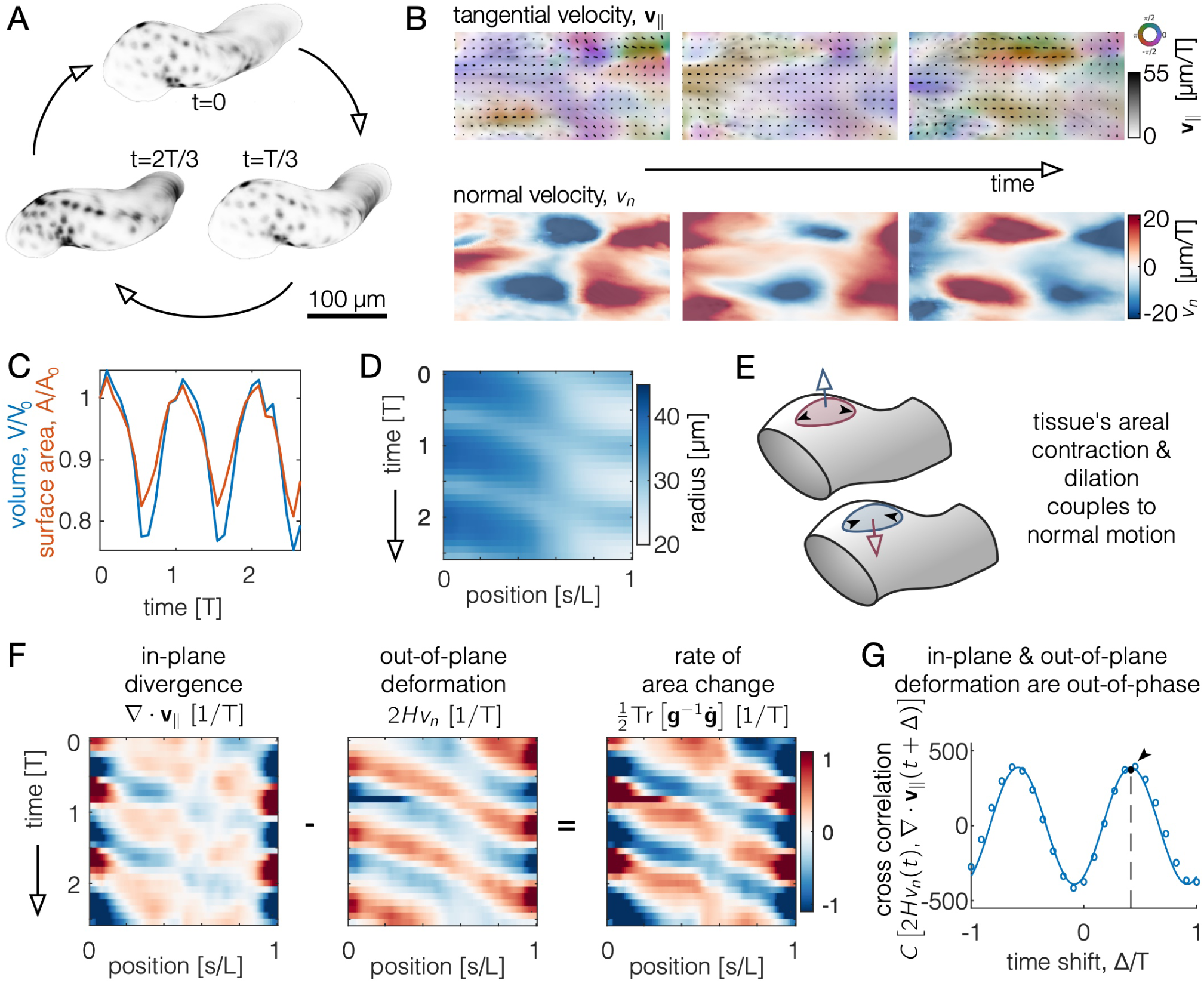
Tracing the in-plane and out-of-plane dynamics of a beating heart reveals the phased coupling between dilatational and transverse deformations. *(A)* Segmentation of a beating zebrafish heart shows cyclic deformations in 3D, shown at three equally spaced timepoints within a beat period, *T*. *(B)* Decomposing tissue motion into in-plane and out-of-plane demonstrates how pulsed deformation travels along the long axis of the tube. The tangential velocity is represented as a color denoting its direction along the long axis (purple or orange) or along the circumferential axis (green or pink), with magnitude denoted by opacity. Inward motion corresponds to *v*_*n*_ *>* 0. *(C)* Both the total enclosed volume and the surface area oscillate over time. *(D)* A kymograph of the radius of the tube measured along the long axis shows cyclic beating. We average the radius around the circumferential axis for each axial position *s* in this plot. *(E)* During each cycle, tissue undergoes both out-of-plane motion and in-plane deformation. These two are coupled, such that the rate of area change depends on both the normal motion and the divergence of the in-plane velocity. *F)* Kymographs of in-plane and out-of-plane motion averaged along the circumferential axis highlights waves of contraction. During each cycle, the in-plane and out-of-plane deformations are nearly out of phase, so that the rate of local tissue area change is large. *(G)* Cross correlation between in-plane tissue dilatation and out-of-plane deformation (constriction of the heart tube) indicates an offset phase relationship. The curve shown is a fit to the data by an offset sinusoidal wave, with a peak at Δ = 0.416 ± 0.006 *T*.

### Beating heart exhibits a phase delay between in-plane motion and out-of-plane deformation

Cyclic deformations of the heart result in in-plane velocities **v**_∥_, colored by their orientation and out-of-plane motion *v*_*n*_ that constricts or dilates the tube (in Fig. 4B). The extracted heart shape changes both surface area and volume as it beats, shown in Fig. 4C. Fig. 4D highlights the waveform of the beat by plotting a kymograph of the radius as a function of time and position along the long axis of the tube. In this measurement, we average along the circumferential axis of the tube, given that the developing heart is reasonably symmetric along its circumference at this stage.

Unlike in midgut morphogenesis, the beating heart’s in-plane velocities are not proportional to the out-of-plane deformation so as to produce incompressible motion (Fig. 4E-G). While both the in-plane divergence and out-of-plane motion display directional waves in their kymographs, the two fields are nearly – but not fully – out of phase. As shown in Fig. 4G, we measure a phase offset between the two fields of 0.416 ± 0.006 *T*, where *T* is the period of heart beating. In other words, as the tube constricts in the normal direction, the tangential velocities are compressive, such that the tissue locally changes area in an oscillating manner. This feature contrasts sharply with the irreversible constrictions of the fly midgut during embryonic stages 15-16, in which nearly incompressible kinematics lead to a 97% correlation between the two fields, resulting in small but persistent areal growth in the lobes and slight areal contraction near the constrictions (Fig. 3E and [4]). This analysis offers a route for quantitatively testing mechanical models of the beating heart.

### Mode decomposition of the heart reveals two out-of-phase characteristic deformations

Finally, we applied TubULAR’s mode decomposition tools to explain the complex cyclic beating of the embryonic zebrafish heart by simpler constituent motions. To do so, we performed principal component analysis (PCA) on the set of 3D tissue velocities across the surface over time. This determines the motions in the material coordinates that explain the majority of the variance in the data over time. The most prominent modes are displayed in Fig. 5A-B. TubULAR performs a Helmholtz-Hodge decomposition of these modes using DEC to probe the signatures of motion contributing to each mode.

**FIG. 5.**
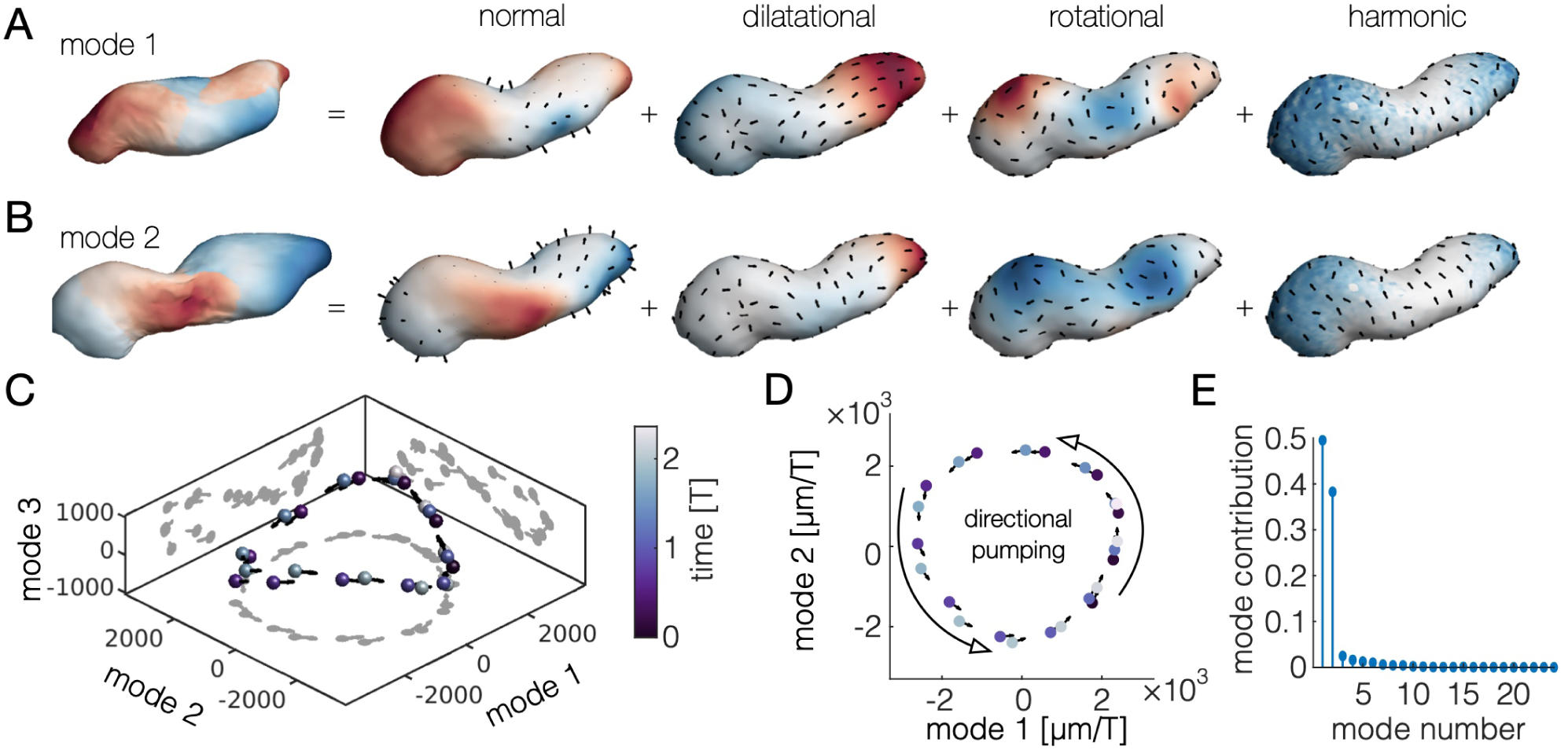
Decomposition of tissue motion in the beating heart reveals two principal components with non-reciprocal dynamics. *(A-B)* Visualization of the principal components of Lagrangian tissue velocities. The first image in each panel shows the deformation induced by moving along the associated component axis, colored by the alignment of the component axis with the surface normal direction. Subsequent images illustrate the Helmholtz-Hodge decomposition of the tangent part of the mode and also the normal part of the mode. Color in the dilatational part, rotational part, and harmonic part are given by the scalar potential, vector potential, and norm of the harmonic mode respectively. Color in the normal part is given by the norm of the normal component of the mode. *(C-D)* Principal component analysis on the time course of deformations returns two dominant modes of deformation with a phase relationship of nearly *π/*2, such that the system winds in state space along a nearly planar, circular pattern. *(E)* A comparison of the relative contribution of the first 24 modes. Mode contribution is defined as the time averaged ratio of the squared length of the projection of the velocity along each mode normalized by the total squared length of each velocity vector in state space.

We find that two modes dominate the dynamics, offering insight into the kinematics driving unidirectional pumping. As shown in Fig. 5C-D, the system oscillates between the first two modes, sweeping out a roughly circular trajectory subtending a nonzero area. Other pairings, in contrast, generate trajectories that subtend approximately zero area (Fig. 5C and Supplementary Information). This phased oscillation between the first two principal components indicate a simple description underlies unidirectional pumping of the heart. Computing the contribution of each mode to the total motion validates this 2D state representation: the first two modes capture nearly 90% of the deformation (Fig. 5E).

## DISCUSSION

We have developed a computational framework for unravelling the complex, dynamic shapes of tube-like surfaces into their principal signatures of deformation, providing a documented, open-source MATLAB implementation to the community. Our implementation unifies the core elements of this toolkit with the existing ImSAnE package [24]. This framework computes Lagrangian measures of strain and strain rate, decomposing dilatational and rotational signatures and mapping them onto the a reference material configuration. Using this approach, we characterized the tissue dynamics and strains during midgut morphogenesis. We then characterized the cyclic deformations of the beating zebrafish heart – highlighting a out-of-phase relationship between in-plane motion and out-of-plane deformation – and captured the heart’s directional pumping motion with a two-dimensional principal component analysis.

An efficient method for tracing surface dynamics in the Lagrangian frame of reference offers new opportunities for understanding not only organ dynamics during morphogenesis, but also organoid systems and sub-cellular structures. To illustrate this, we captured a surface representation of a developing neural tube derived from human stem-cells [55] and tracked a deforming phase-separated droplet in a microtubule gel [56], each shown in the Supplementary Information.

A remaining challenge is to extend methods of tracking deformation through branching events and changes in the connectivity of the surface. Here we addressed the challenges of complex and dynamic geometries and are even able to capture topological changes in the winding of tube-like surfaces. Yet, we focused our efforts to follow tubes with a single opening on each end. Extending to higher-order networks of tubes [57] and shapes which fuse or separate [55] poses future challenges. As multi-scale datasets emerge, we forsee constrained parameterization methods as useful building blocks for tracking the dynamics of hierarchical processes such as the branching lung or kidney [58].

## ACKNOWLEDGEMENTS

Sebastian Streichan and Boris Shraiman provided crucial insights, discussions, mentorship, and expertise. Sebastian Streichan additionally provided the laboratory and computational resources to develop and execute this work, with primary support for this work from NSF Grant No. PHY-2047140. We thank Sebastian Streichan for the light sheet dataset of the beating zebrafish heart, Eyal Karzbrun and Sebastian Streichan for the confocal dataset of an *in vitro* neural tube used in the Supplementary Information, and Alexandra Tayar for the dataset of the deforming DNA droplet in a microtubule gel used in the Supplementary Information. We also thank Suraj Shankar for useful discussions. NPM acknowledges support from the Helen Hay Whitney Foundation. DJC acknowledges support from the NSF Grant No. PHY-1707973. The work was also supported in part by the National Science Foundation Grant No. NSF PHY-1748958.

## Notes

https://npmitchell.github.io/tubular/

### Competing Interest Statement

The authors have declared no competing interest.

### Summary of Updates

Scope and clarity of presentation, added Supplementary Information

