## Supplementary Information for "TubULAR: Tracking *in toto* deformations of dynamic tissues via constrained maps"

Noah P. Mitchell[1,2,*] and Dillon J. Cislo[2,†]
[1]*Kavli Institute for Theoretical Physics, University of California Santa Barbara*
[2]*Department of Physics, University of California Santa Barbara*

### CONTENTS

Overview of approach by example 1

Surface extraction using level sets 1
Segmentation of the zebrafish heart 2

Constrained mapping to the plane follows tissue motion 2
Initial conformal map $f$ 3
Ricci flow 3
Annular orbifold map 4
Independence of mapping on choice of longitudinal seam 5
Quasiconformal map $\Phi \circ Z$ to $(s, \phi)$ coordinates 6
Auxiliary geometric descriptors of surface dynamics: centerline and effective radii 7
Constrained parameterization defines a system-spanning centerline of the surface 7
Constrained parameterization defines an effective radius along the surface 7
Refined Lagrangian parameterization of the surface, $\varphi = J \circ \Phi \circ Z \circ f$ 7
Surface velocities and discrete exterior calculus 8
Helmholtz-Hodge decomposition of vector fields on dynamic surfaces 10
Lagrangian measures of time-integrated tissue strain 10

Integration of TubULAR with ImSAnE 10
Passing ImSAnE to TubULAR 10
Upgrades to ImSAnE integrate TubULAR's functionality 10

Example of inferring intercalation rates ('tissue tectonics') using TubULAR 11

Analysis of beating zebrafish heart dynamics 12

A cytoskeletal gel actively deforms liquid droplets 12

References 13

### OVERVIEW OF APPROACH BY EXAMPLE

In addition to detailed documentation and example pipelines available on GitHub, we summarize the steps in our approach in Fig. S2. This gives a typical sequence of method calls to extract surfaces, create an initial sequence of parameterizations constrained for minimal tissue motion in the pullback plane, and compute covariant measures of tissue dynamics. Subsequent steps further refine the material coordinate definition to remove residual motion and read out measures of motion and strain in this Lagrangian frame (along 'material pathlines'). The last set of steps visualize the material motions, decompose them into divergence, curl, rate of area change, and measures of anisotropic deformation like the Beltrami coefficient. Finally, methods compute principal components of the tissue dynamics and decompose into eigenfunctions of the Laplace-Beltrami operator, akin to spherical harmonics for a sphere or Bessel functions for a cylinder but defined on an arbitrary surface. Fig. S3, meanwhile, gives an overview of the class structures included in the toolkit.

### SURFACE EXTRACTION USING LEVEL SETS

To extract whole-organ surfaces, we use a level sets approach, combined with marching cubes [2] and Laplacian smoothing. While the literature on level sets segmentation is vast [3], we give a brief overview of the relevant method here.

The process of surface detection is mapped onto an optimization problem by defining a physics-inspired cost functional [4, 5]:

$$F[c_1, c_2, \mathcal{S}] = \mu \int_{\mathcal{S}} \mathrm{d}s + \nu \int_{\Omega} \mathrm{d}^3\mathbf{x} \tag{1}$$

$$+ \lambda_1 \int_{u>0} |I(\mathbf{x}) - c_1| \mathrm{d}\mathbf{x} + \lambda_2 \int_{u<0} |I(\mathbf{x}) - c_2| \mathrm{d}\mathbf{x}, \tag{2}$$

where $c_1$ and $c_2$ are the average values of the data inside and outside, respectively:

$$c_1(\mathcal{S}) = \langle I(\mathbf{x}) \rangle_{\text{inside}} \tag{3}$$

$$c_2(\mathcal{S}) = \langle I(\mathbf{x}) \rangle_{\text{outside}}. \tag{4}$$

The first term is a surface tension that tries to smooth out ruggedness in the surface. The second term is an effective pressure that penalizes blow up in the enclosed volume of the segmented regions. The final two attachment terms incorporate the actual measured data into the optimization procedure. The third term attempts to homogenize the intensities of voxels included in the

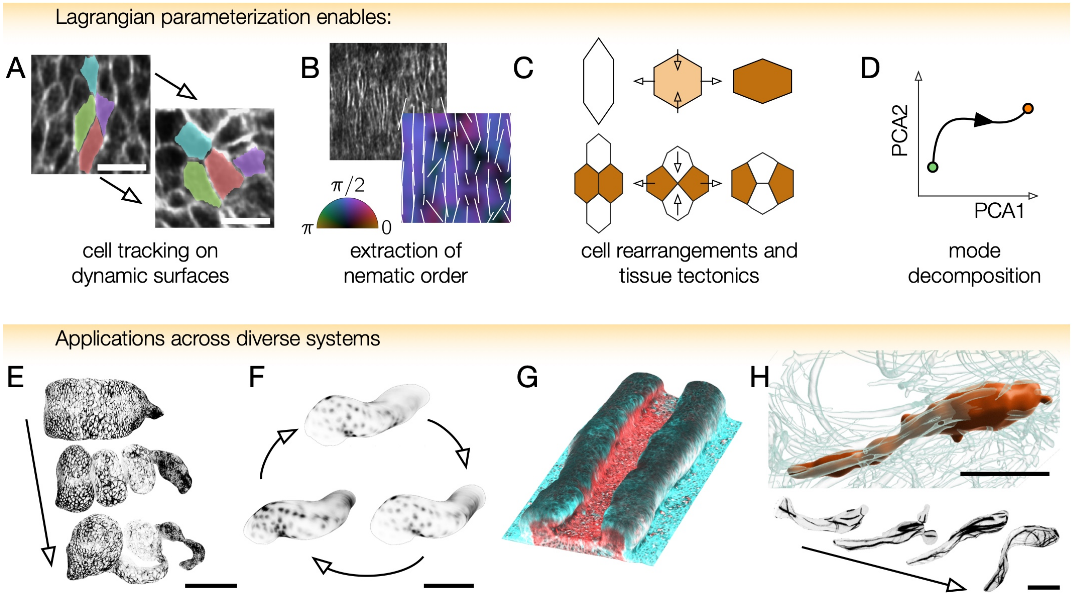


FIG. S1. *(A)* Computational microscopy enables dynamic tissue surface extraction and cell tracking. After selectively imaging the endodermal layer in a *w;48Y-GAL4/UAS-CAAX:mCherry;klar* embryo, we track cells, with four highlighted that exhibit intercalations highlighted here. Scale bars are 10 $\mu$m. *(B)* Performing a radon transform on a patch of circular visceral muscle cells – which form a 'palisade' structure ensheathing the midgut endoderm – returns a measure of tissue anisotropy aligned with the circumferential direction ($\phi = \pi/2$, violet). *(C)* Constructing material-frame pullback images facilitates measurements of tissue shape change, cell shape change, and cell intercalation rates (see Supplementary Information). *(D)* Following the tissue's deformation enables mode decomposition of the dynamics, offering descriptions with reduced complexity, shown schematically as a path of the system through mode space after performing principal component analysis (PCA). *(E-H)* This method tracks deformations for organs and *in vitro* systems alike, including the Drosophila midgut, the beating zebrafish heart, a stem-cell-based neural tube, and a phase-separated droplet deforming in a cytoskeletal gel. Scale bars are 100 $\mu$m for all panels. *(G)* Characterizing the tissue surface offers a route to study neural tube development [1]. *(H)* We follow a DNA-nanostar-based droplet (red) deforming in an active fluid. Here, mechanical coupling between the interface of a liquid-liquid phase separated DNA droplet and a surrounding active microtubule fluid (cyan rods) generates continuous deformation towards droplet breakup. In the time series, microtubule fluorescence within 10 $\mu$m of the interface is projected onto the deforming droplet surface.

interior of the segmented volume and the fourth term attempts to homogenize the intensities of the excluded voxels.

To segment the midgut and heart surfaces, we generate a level set solution for a contiguous volume enclosing the interior of the tube, partitioning space into a topological sphere (the 'filled' tube) and its exterior. TubULAR then removes endcaps of the mesh at the interface to create a topological tube (see later discussion).

#### Segmentation of the zebrafish heart

The zebrafish heart data set posed particular challenges requiring the application of some non-standard segmentation procedures, which we summarize here. Rather than attempt to segment both the heart tissue and the space it enclosed, we used the level set methods to segment only the heart tissue. This resulted in a binary level set solution of toroidal topology. We then applied a homotopic thinning procedure slice-by-slice along the length of the tube to produce a point cloud approximating the mid-surface of the heart tissue. We then fed a smoothed, up-sampled version of this point cloud into our Poisson surface reconstruction algorithm to produce a closed, sphere-like mesh of the heart. The results at this point in the segmentation process were structurally identical to those produced in a typical pipeline.

### CONSTRAINED MAPPING TO THE PLANE FOLLOWS TISSUE MOTION

To follow tissue surfaces as they deform, we begin with an initial map at a reference timepoint that defines the material coordinates, then construct maps to minimize subsequent tissue motion in the pullback plane, remove any residual motion, and generate material pathlines in 3D. We denote the dynamic map from the evolving surface to a fixed 2D material coordinate system as $\varphi(t)$. This dynamic map to a fixed material coordinate system is built via a sequence of four steps: $\varphi(t) \equiv J \circ \Phi \circ Z \circ f$, each of which is detailed in this section. Briefly, $f : \mathcal{S}(\mathbf{x}) \to (u, v)$ is a conformal map of

***Instantiation***
Set experiment metadata and any custom options..........................xp=struct(...); opts=struct(...);
Instantiate TubULAR....................................................tubi = TubULAR(xp, opts)

***Surface extraction***
Create downsampled volumes to use for iLastik & level sets.............tubi.prepareIlastik()
Extract the surfaces using level sets on output of iLastik or raw data..tubi.getMeshes()

***Parameterization***
Define global reference frame (APDV)...................................tubi.computeAPDVCoords()
Define endcap points and one point for a virtual seam..................tubi.computeAPDpoints()
Align meshes into global frame.........................................tubi.alignMeshesAPDV()
Optional: render data on dynamic surfaces (can be slow)................tubi.plotSeriesOnSurfaceTexturePatch()
Compute initial centerlines............................................tubi.extractCenterlineSeries()
Temporally average centerlines and fix any inconsistencies.............tubi.generateCleanCntrlines()
Remove the endcaps.....................................................tubi.sliceMeshEndcaps()
Clean up the mesh endcaps..............................................tubi.cleanCylMeshes()
Cut the meshes along virtual seams.....................................tubi.generateCurrentCutMesh()
Constrained parameterization of the mesh into (u,v) and (s,$\phi$) frames....tubi.generateCurrentSPCutMesh()
Generate pullback images...............................................tubi.generateCurrentPullbacks()

***Dynamics***
Initial pass of smoothing embedding over time..........................tubi.smoothDynamicSPhiMeshes()
Create pullbacks of smoothed (s,$\phi$) frames..............................tubi.generateCurrentPullbacks()
Tile the images in $\phi$ (as 'double-cover')...............................tubi.doubleCoverPullbackImages()
Measure any residual material motion in the parameterization...........tubi.measurePIV2d()
Translate pullback information into motion in 3D (embedding space)......tubi.measurePIV3d()

***Refined processing of surface dynamics***
Recommended: smooth the resulting 3D dynamics along material pathlines..tubi.timeAverageVelocities()
Compute material pathlines in 3D.......................................tubi.measurePullbackPathlines()
Measure rate of strain over the surface at each timepoint..............tubi.measureStrainRate()
Query the rate of strain along pathlines...............................tubi.measurePathlineStrainRate()
Measure the integrated strain along pathlines..........................tubi.measurePathlineStrain()

***Interpretation and mode decomposition***
Visualize flows........................................................tubi.plotTimeAvgVelocities()
Decompose into divergence and curl, scalar potential fields............tubi.helmholtzHodge()
Compute areal rate of change...........................................tubi.measureMetricKinematics()
Mesaure anisotropic strain using Beltrami coefficient..................tubi.measureBeltramiCoefficient()
Decompose into Principal Component Analysis............................tubi.getPCAoverTime()
Decompose into eigenmodes of the Laplace-Beltrami operator.............tubi.getLBSoverTime()

FIG. S2. **Example high-level pipeline for data analysis using TubULAR passes through constrained parameterization, measurement of surface dynamics, refinement, and steps to interpret the results.** Descriptions of typical steps in a TubULAR pipeline are listed on the left, with the corresponding class method calls for each goal on the right.

the surface $\mathcal{S}$ to the unit square (via Ricci flow or Dirichlet energy minimization). $Z : u \to s$ maps each longitudinal coordinate $u(t)$ to proper length $s(u(t))$ along the long axis of the tube-like surface. $\Phi : v \to \phi$ stabilizes motion of the tissue along the circumferential axis. Finally, $J : (s, \phi) \to (s_0, \phi_0)$ removes any residual motion of the material in the pullback plane. Let us turn to each component in turn.

### Initial conformal map $f$

We homotopically flatten the 3D surface to the plane using one of two methods. The first (default) option is Ricci flow, which results in a precisely conformal output at the cost of being slow, while the second option is using a map minimizing a Dirichlet energy, which in general is faster, but produces less conformal results.

#### *Ricci flow*

Originally introduced by Hamilton in the context of geometric topology, Ricci flow is a tool that enables the design of Riemannian metrics with prescribed curvatures. Ricci flow deforms a Riemannian metric proportionally to its intrinsic curvature, such that the curvature evolves according to a nonlinear heat diffusion process and eventually becomes constant everywhere. In the continuous setting, Ricci flow on 2D surfaces can be defined as

$$\frac{dg_{ij}(t)}{dt} = -2(K(t) - \bar{K})\, g_{ij}(t), \tag{5}$$

where $g_{ij}(t)$ is the time dependent metric of the surface, $K(t)$ is the associated Gaussian curvature, and $\bar{K}$ is the target Gaussian curvature. It is immediately apparent from Eq. (5) that surface Ricci flow is conformal, i.e. preserves angles defined by $g_{ij}(t)$. It has also been demonstrated that the Gaussian curvature during the flow always remains bounded.

Surface Ricci flow has intuitive geometric interpretations, which directly inform the design of data structures in the discrete setting. For instance, the surface can be represented as a mesh triangulation and the metric tensor can be simply represented as a set of positive edge lengths on this triangulation satisfying the triangle inequality. Crucially, it is possible to reformulate discrete surface Ricci flow as a convex optimization problem over the space of discrete metrics, which has a unique minimum and can be solved efficiently using Newton's method. Intuitively, given an initial metric, the method first constructs a circle-packing metric, i.e. it instanti-

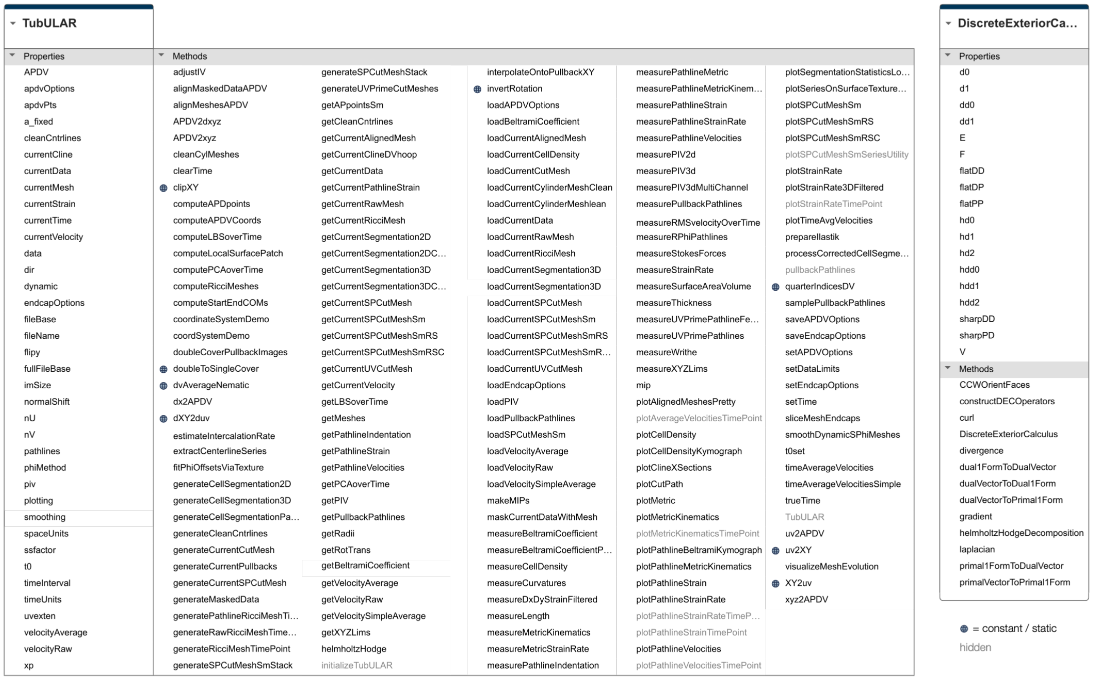


FIG. S3. **The TubULAR class and the DiscreteExteriorCalculus class within `DECLab` carry properties and methods, listed here for reference.**

ates a set of circles living at mesh vertices with consistent conditions on how the circles intersect. Each variational step in the minimization process flows the discrete metric closer to the target curvature in a way that always maintains the integrity of the circle packing. The conformality of the discrete flow is ensured by construction since it always maps (discrete) infinitesimal circles into (discrete) infinitesimal circles. For a comprehensive review of the method, please consult [6].

*Annular orbifold map*

The annular orbifold map follows the formalism of Aigerman & Lipman [7], but applies these ideas to a topological cylinder mapped to a rectangle that is periodic along one dimension. Essentially, we produce a parameterization that minimizes the Dirichlet energy (see next section), enforcing certain boundary conditions so that the resulting parameterization is both seamless and globally bijective.

For a parameterization $\vec{u} = (u, v)$, the Dirichlet energy is defined as

$$E_D = \frac{1}{2} \int ||\nabla \vec{u}||^2 \mathrm{d}A. \tag{6}$$

This energy penalizes ruggedness in $\vec{u}$ and, as a result, its minimizers are smooth, well-behaved parameterizations. Extrema of this energy functional can be found according to a variational principle by solving the associated Euler-Lagrange equation:

$$\nabla^2 \vec{u} = 0. \tag{7}$$

This is, of course, simply the vector Laplace equation. The associated solutions are called *harmonic maps*. In practice, since $\vec{u}$ is unknown, the operator $\nabla^2$ is taken to be the Laplace-Beltrami operator defined on the input 3D surface. Additionally, we demand that the anterior and posterior endcaps of the cylindrical mesh are mapped to vertical lines $u = 0$ and $u = 1$. We also enforce the orbifold condition that corresponding vertex pairs on each cut path map to the same location in the planar parameterization space, modulo a vertical offset corresponding to the periodic spacing along the azimuthal direction of the tube. These boundary constraints ensure that the planar parameterization retains the cylindrical topology of the 3D surface under proper tiling of the plane. This entire problem, including the constraints, can be cast as a single linear system

$$A\vec{u} = b. \tag{8}$$

Any conformal parameterization of the surface will be a minimizer of the Dirichlet energy. To see this, let us specialize for a moment to the case of planar transformations $f : \mathbb{C} \to \mathbb{C}$. Let $z = x + i\,y$ denote a complex coordinate defined over the input domain being parameterized (the analogue of the input 3D surface) and let

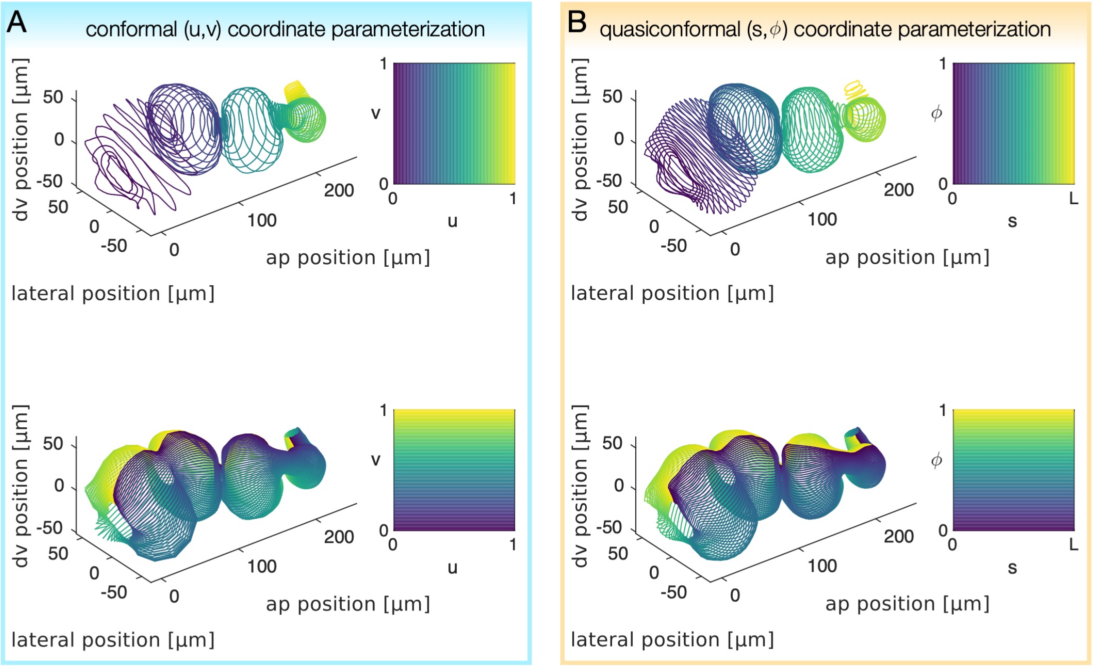


FIG. S4. **A quasiconformal mapping from $(u, v)$ to $(s, \phi)$ aids in spatially uniform surface sampling and velocimetry for refined tissue tracking.** *(A)* The constrained conformal map to the plane at a time 90 minutes after the onset of constrictions ($t = 90$ min) demonstrates large variations in the sampling density along the longitudinal direction (top). Note the high density of circumferential hoops near constrictions and near the posterior tip. *(B)* After creating the conformal mapping to $(u, v)$, the TubULAR workflow constructs an second map to a parameterization in which the surface is more evenly sampled along the longitudinal axis and in which motion in the $v$ direction is subtracted off. Note the more uniform longitudinal density of circumferential hoops (top) and the difference in the curve $\phi = 0$ at this later timepoint, which matches the previous position of the material along $\phi = 0$ at earlier timepoints. At time $t = t_0 = 0$ min, $\phi = v$, while at this later timepoint, $\phi \neq v$.

$f = u + i\,v$ be a complex representation of the parameterization. A conformal parameterization must satisfy the Cauchy-Riemann condition

$$\partial_{\bar{z}} f = 0. \quad (9)$$

It is therefore clear that any conformal mapping must also be harmonic since $\nabla^2 f = 4\partial_z \partial_{\bar{z}} f = 0$. Similar generalized arguments prove the harmonicity of 2D conformal parameterizations of 3D surfaces. In practice, the boundary conditions we enforce preclude the possibility of a truly conformal mapping. However, the results are generally a good approximation of a discrete conformal mapping, especially in the bulk away from the mesh boundaries. The high quality of Dirichlet energy minima as approximations to discrete conformal mappings has also been observed in a variety of settings [7]. While such methods are not currently implemented in TubULAR, we note that the conformality of the parameterizations can be improved by replacing the Dirichlet energy with the so-called *conformal energy*, $E_C = E_D - A(\vec{u})$, where $A(\vec{u})$ denotes the area of the domain of parameterization [8]. This improvement in conformality comes at the expense of increased computational cost and usually do not preserve angles as well the conformal Ricci maps anyway.

*Independence of mapping on choice of longitudinal seam*

The choice of cut path along the long axis of the organ for unrolling the cylinder should intuitively be immaterial to the position of the mapped surface coordinates. For the Ricci flow case, this is true by construction: we map to an annular domain (with one endcap boundary mapped to the unit circle and one near the origin), then take a logarithm of these coordinates to acquire a rectilinear representation. For the annular orbifold map, we must enforce this path independence.

Inspired by the topology-preserving orbifold mappings of spherical surfaces [7], we enforce a set of boundary conditions to ensure that the parameterization in the plane respects the cylindrical topology of the 3D surface. Namely, we demand continuity in the boundary components of the 2D meshes associated with the virtual seam, such that, if one were to tile these meshes in the plane, moving across this boundary from one tile into another would be physically indistinguishable from crossing over the virtual seam on the 3D cylindrical surface. This mapping ensures that the edges composing the cut path takes on a unique shape in the domain of parameterization, and the shape of these edges would be identical whether or not the edges are components of the cut path.

There is one caveat to this independence: the cut path will change the output pullback map if it winds around the tube relative to the centerline with a different winding number. Therefore, we take measures to enforce this topological constraint. For the first timepoint mapped to the plane ($t_0$), we choose a cut path to be the geodesic connecting the two endcaps. For subsequent (and previous) timepoints, we ensure the winding number does not change relative to this path's winding.

While several options are available for enforcing this constraint, the default behavior uses a crude approximation to the centerline and measures the winding of the cut path around this curve. The details of this centerline construction are not particularly important, since the curve's purpose is to provide a topological constraint – not a geometric one – on the cut path chosen, whose own geometry is immaterial so long as it does not wind around the centerline. Nevertheless, we give a brief description here: Briefly, for each timepoint, we measure a crude pathline via fast marching (Fig. S6B). We connect the endpoints by a curve that spans the interior of the mesh found by minimizing the 'time of travel' with a speed of travel through any given voxel determined by the distance transform of the segmented data volume [9]. If a geodesic path is found to change its winding from one timepoint to the next, the path is perturbed to more closely match the previous one in space until the topology is preserved. We found this 'trial and error' approach to be far faster than constructing an explicitly-topologically-equivalent curve (for example, after Ricci flow to an annular domain). The relevant TubULAR methods provide options for choosing different approaches if needed, including the explicit construction method.

#### Quasiconformal map $\Phi \circ Z$ to $(s, \phi)$ coordinates

We then introduce a further coordinate transformation which we found aids in surface stabilization. Because directly measuring optical flow in $(u, v)$ coordinates is often too crude and does not uniformly sample tissue motion across the embedding surface, we followed a more constrained approach to remove motion, which we found to aid in subsequent refinement of the tissue stabilization. We denote this second planar parameterization $(s, \phi)$, shown in Fig. S4B. We find this $(s, \phi, t)$ parameterization aids in both visualization and enables more accurate velocimetry measurements than other choices we considered, particularly when large variations appear in the effective radius of the surface along its long axis. This second map, which we denote $\Phi \circ Z$ is a quasi-conformal transformation (i.e. a smooth transformation with finite anisotropic distortion [10]) of the initial $(u, v)$ coordinates.

For the reference timepoint $t_0$ considered first, the coordinate directions $\hat{s}$ and $\hat{\phi}$ are the same as the conformal mapping to the plane at a reference time $t = 0$. Furthermore, at this initial timepoint, $\phi$ is identical to the intrinsic circumferential axis of the conformal map. The sole difference is that $s$ parameterizes a longitudinal position along the long axis of the organ at $t = 0$. In particular, we compute $s$ as the average geodesic length along the surface from the anterior endcap to a set of uniformly-sampled points with fixed horizontal coordinate in the conformal pullback space (Fig. 2A-D). Intuitively then, $s$ this is the average path length required to travel on the surface from the anterior face to a given location along curves of constant $\phi$.

In more detail, we define the $(s, \phi)$ domain of parameterization, which is less conformal but which more equally represents different patches of tissue that initially experience vastly different dilation in the map $f$ from 3D embedding $\vec{x}$ to 2D pullback $\vec{u} = (u, v)$. This empirically improves measurements of tissue velocity in plane for our shapes, and we expect the additional transformation will improve other tissues that are elongated in quasi-axisymmetric geometries. Circumferential 'hoops' of tissue surrounding the centerline that are equally sampled *along* the centerline will be equally spaced in the pullback coordinates. The map $\Phi \circ Z$ from the previous (conformal) frame $(u, v)$ to the $(s, \phi)$ coordinate system is defined by

$$s(u) = \int_0^u \left\langle \frac{\mathrm{d}s}{\mathrm{d}u'} \right\rangle_{u'=\mathrm{const}} \mathrm{d}u', \tag{10}$$

and

$$\phi(u, v) = v - \phi_0(u). \tag{11}$$

Equation 10 ensures that circumferential hoops sampled at equally spaced distances (as measured by their average proper distance) along the longitudinal axis are equally spaced in pullback space.

Equation 11 removes tissue motion along $\phi$ at each longitudinal position $s$. The form of $\phi_0(u)$ is such that motion of the tissue along each hoop is cancelled out in the pullback to acheive a more Lagrangian parameterization. For $t = t_0$, $\phi_0(u) = 0$. For other timepoints, $\phi_0(u)$ is chosen to minimize the difference in positions of material points at the current timepoint relative to the previous (next) timepoint for $t > t_0$ ($t < t_0$). To minimize the difference in material positions, we maximize the correlation of circumferential 'hoops' defined by $s_i < s < s_j$ to the mapped positions at a previously-solved timepoint closer to $t_0$. These hoops can be visualized in pullback space as vertical strips in the $(s, \phi)$ pullback coordinates. If optical flow is well constrained, the user may toggle the option for stabilization method so that $\phi_0(u)$ is defined by maximizing the cross correlation between the intensity data lying within each circumferential hoop and the intensity data in the previous pullback lying near the same

$u$ coordinate. This option uses phase correlation of the pullback image itself and compares each strip in pullback space (corresponding to a hoop in 3D data space) to the previous image. If optical flow is an unreliable measure in this step, we have found that shifting these slices along $\phi$ (which is a periodic dimension) to minimize the difference in 3D space defines $\phi_0(u)$ gives satisfactory results, and further processing in the next stages of the pipeline remove any residual material motion in the $(s, \phi)$ plane.

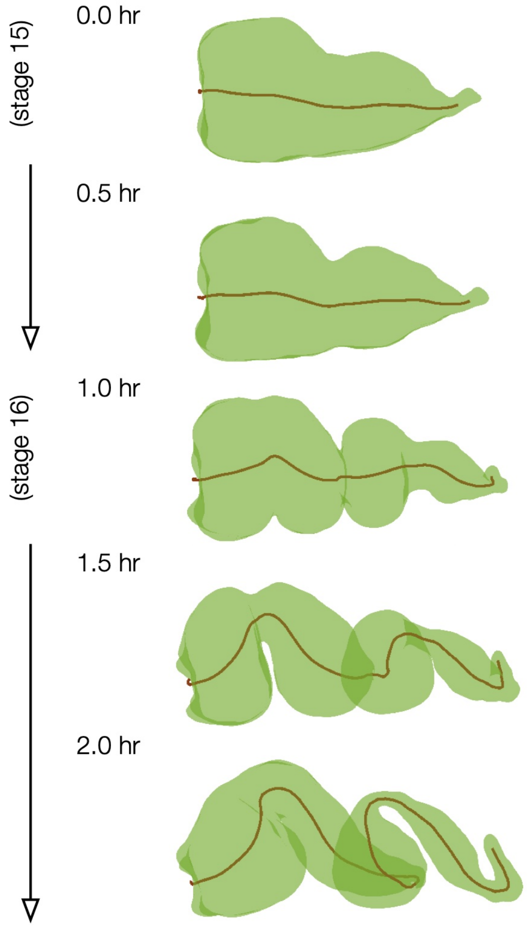


FIG. S5. **Example centerlines computed via TubULAR span the system size and capture a 1D representation of the organ dynamics.** These centerlines follow directly from the material parameterization, with each curve (brown) composed of sequentially-connected centroids of circumferential 'hoops'.

#### Auxiliary geometric descriptors of surface dynamics: centerline and effective radii

In addition to following tissue motion, our parameterization scheme naturally produces a unique route to compute a useful version of the organ's centerline. This definition of centerline has several advantages, including the built-in association of each centerline point with a set of points on the organ surface (a set of mesh vertices) that span a circumferential 'hoop'. Because of this association, the notion of an 'effective radius' at each point along the centerline naturally follows, as shown in Fig. S6.

##### *Constrained parameterization defines a system-spanning centerline of the surface*

Centerline construction leverages the surface parameterization in 3D space already created from the previous step. Hoops for which $s =$ constant define an effective circumference for increments along the length of the organ, and the average 3D position of each hoop defines its centerline point. Connecting mean points of adjacent hoops along the length of the organ defines the centerline of the object. This construction offers several advantages to previous methods of centerline construction.

##### *Constrained parameterization defines an effective radius along the surface*

We define an effective radius as the average distance from each point in a uniform sampling of a ring of constant $s$ (computed via the mapping to the pullback plane) to the centerline, which is composed of the mean positions of all circumferential rings. We then identified constriction locations as rings of constant $s$ whose effective radii $r(s)$ are local minima (so that $\partial_s r(s) = 0$). Local minima in effective radius are tracked starting at the onset of folding forward in time to define constriction locations. Before the onset of folding, presumptive constriction locations are inferred by back-tracing the onset location to earlier timepoints.

We note that this measurement of effective radius could be done on the $(u, v)$ coordinate parameterization and would give identical results, since curves of constant $u$ are also curves of constant $s$. The effective radii $r(s)$ are therefore equal to those indexed by $u$: $r(s(u)) = r(u)$.

#### Refined Lagrangian parameterization of the surface, $\varphi = J \circ \Phi \circ Z \circ f$

After the previous constrained parameterization into the plane via $\Phi \circ Z \circ f$, which minimizes much of the tissue motion in the parameterization plane, we accomplish further refinement via computing pathlines in the domain of parameterization using particle image velocimetry (PIV). Advecting mesh vertices along these pathlines, then inverting the dynamic map to the plane gives the 3D positions of material points as they deform. This provides the surface shown in Fig. 1D of the main text, for instance.

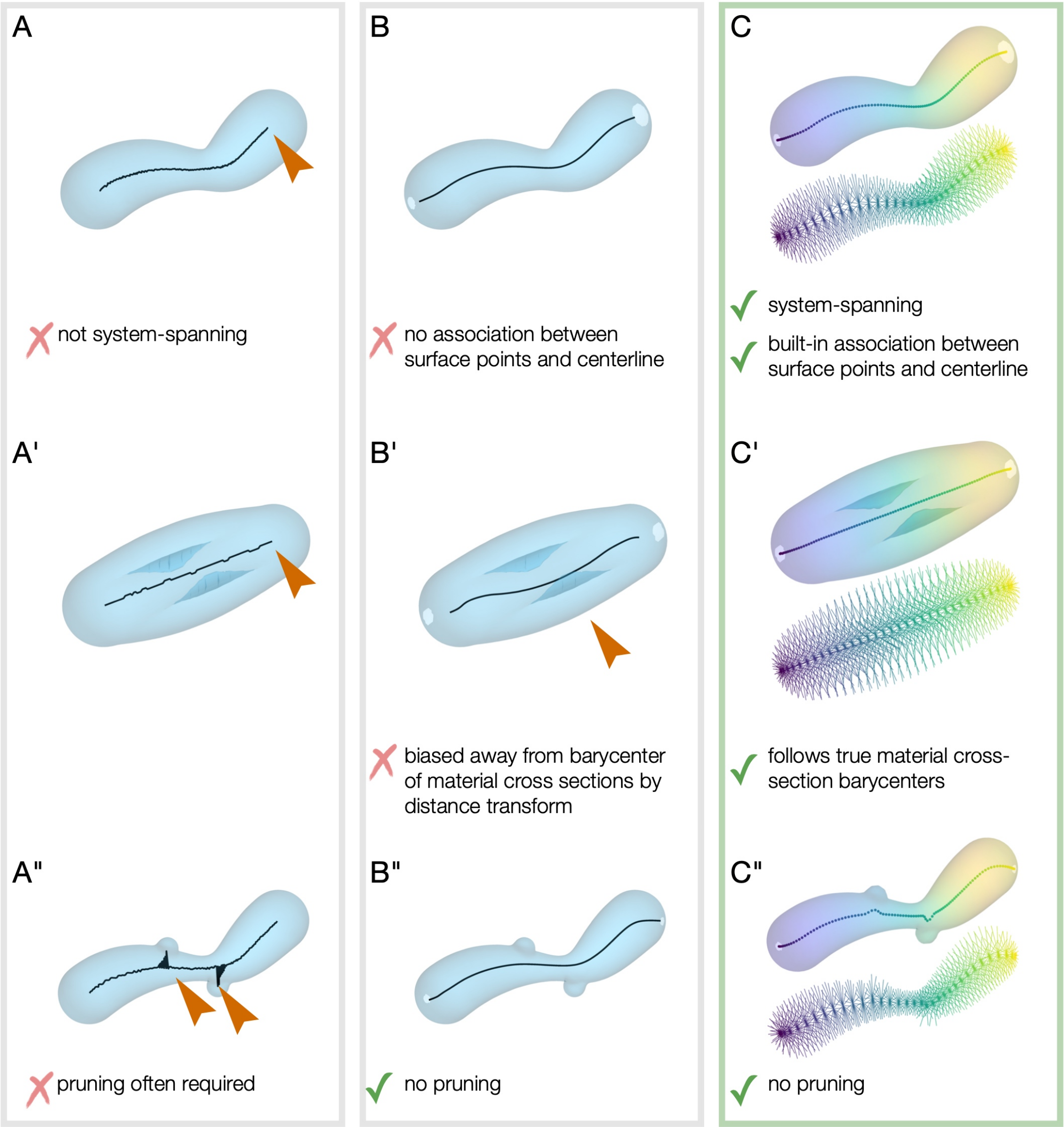


FIG. S6. **Our method provides a geometric route to centerline extraction which offers several benefits over traditional approaches.** *(A-A")* An approximate centerline can be built from homotopic thinning methods [11]. This simple method has several downsides, including that the curves do not span the whole system, and pruning of the curve is needed when the surface is not sufficiently smooth. Associating the mesh surface and the curve poses conceptual challenges as well. *(B-B")* An alternative – which is implemented in an auxiliary step for constraining surface parameterization in TubULAR – minimizes the 'time of travel' within the segmented volume from selected endpoints, where the speed of travel through a given voxel is weighted by its signed distance from the mesh surface [9]. While this offers system-spanning centerline curves, the curves deviate from the center of the object when the surface is puckered, as in panel B'. Additionally, no association between the mesh surface the the curve is given by this method. For contorted tubes such as the fly midgut, nearest distance matching gives spurious associations. *(C-C")* Constructing centerlines based on the coordinate parameterization to the material frame offers advantages for finding a single curve without branching with explicit associations between surface points and the centerline. This construction enables examination of the constriction cross sections in the main text.

#### Surface velocities and discrete exterior calculus

3D velocity vectors arise naturally from our approach via mapping the endpoints of 2D PIV vectors into their respective 3D surfaces. Geometrically, displacement vectors $\mathbf{v}$ extend from one coordinate $\mathbf{x}_0$ in 3D on the surface at time $t_0$ to a different coordinate $\mathbf{x}_1$ on the deformed surface at time $t_1$. When $t_0$ and $t_1$ are adjacent timepoints, this defines the 3D tissue velocity at $t_0$ as $\mathbf{v} = (\mathbf{x}_1 - \mathbf{x}_0)/(t_1 - t_0)$. We then decompose the velocity into a component tangential to the surface $\mathbf{v}_\parallel$ and a normal component $v_n$.

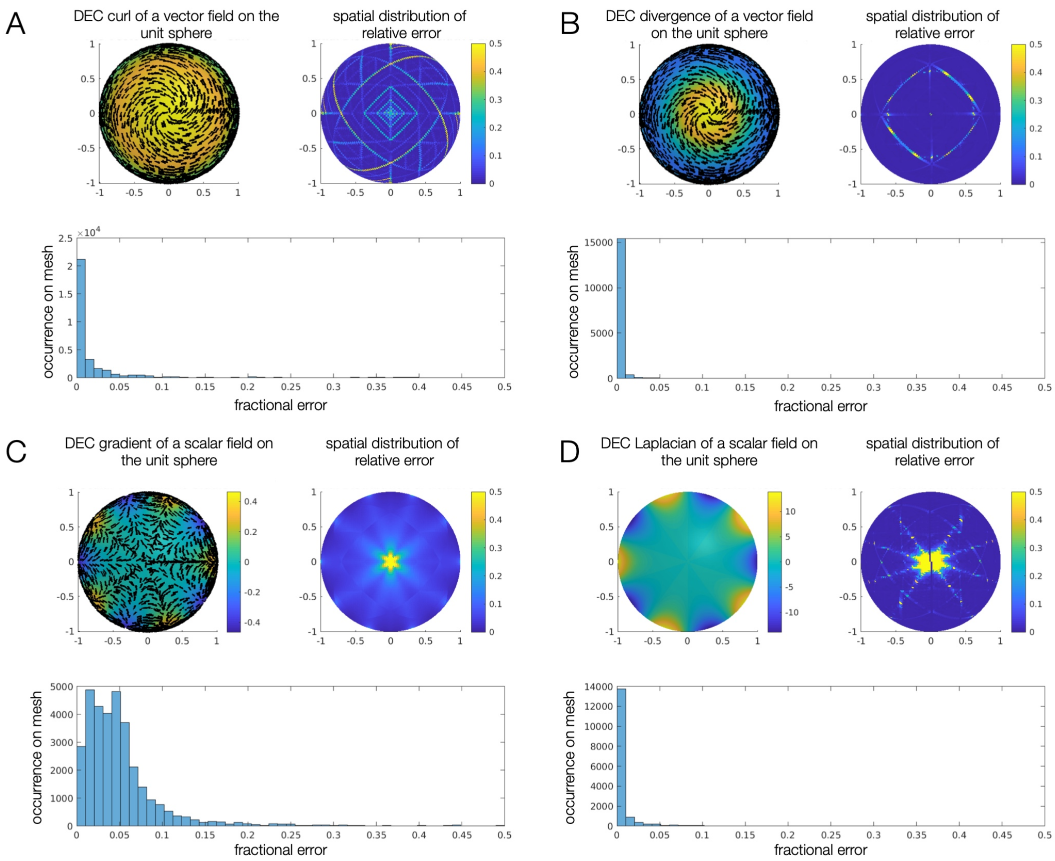


FIG. S7. **DECLab operators reproduce analytic results, with some mesh-dependent errors.** Examples of computed curl $((\star d(\vec{v}^\flat))^\sharp)$, divergence $(\star d \star (\vec{v}^\flat))$, gradient $((d\varphi)^\sharp)$, and Laplacian $(\star d \star d\varphi)$ fields on the surface of a sphere show little relative error, plotted as histograms for each case. Values are compared with analytic results for the tested vector fields $\vec{v}$ in (A) and (B) and scalar fields $\phi$ in (C) and (D). Further details and example code are available on the GitHub documentation page.

The tangential velocity fields $\mathbf{v}_\parallel$ can then be further analyzed using our implementation of the discrete exterior calculus (DEC). DEC discretizes the methods of exterior calculus in the continuous setting for application on simplicial complexes such as mesh triangulations. DEC is built using a straightforward set of discrete differential forms, defined on mesh vertices, edges, and faces. On a 2D surface, the only such forms are 0-forms (scalars), 1-forms (analogues of vector fields), and 2-forms (oriented areas). The DEC also defines representations of the exterior derivative $d$ and the Hodge star $\star$ in terms of simple linear operations. These elemental operations are maps between the different spaces of $k$-forms on the mesh ($k \in \{0, 1, 2\}$). This technology can be exploited for a wide variety of applications in discrete geometry processing. In particular, it allows us to easily compute gradients of the velocity field (or other vector/tensor fields) on curved surfaces.

For a given mesh, instantiating `DECLab`'s `DiscreteExteriorCalculus` class generates the elemental operators, $d$ and $\star$, for each possible pairing of $k$-form types, i.e. for a one form $\omega$ the operation $d\omega = \alpha$ generates a 2-form $\alpha$. When strung together, these operators generate familiar mathematical operations such as the divergence, curl, and Laplacian – except now these operators take care to incorporate the curvature and geometry of the triangulated surface. For completeness, we enumerate some familiar differential operations in the language of exterior calculus. Let $\varphi$ denote a scalar field (0-form), let $\vec{v}$ denote a vector field, and let $\flat/\sharp$ denote the musical isomorphisms that transform vector fields into 1-form fields and 1-form fields into vector fields, respectively. Then, common differential operations in the language of exterior calculus are:

$$\nabla\varphi \to (d\varphi)^\sharp \tag{12a}$$

$$\nabla^2\varphi \to \star d \star d\varphi \tag{12b}$$

$$\nabla \cdot \vec{v} \to \star d \star (\vec{v}^\flat) \tag{12c}$$

$$\nabla \times \vec{v} \to (\star d(\vec{v}^\flat))^\sharp \tag{12d}$$

$$\nabla^2\vec{v} = ((\star d \star d + d \star d\star)\, \vec{v}^\flat)^\sharp. \tag{12e}$$

Fig. S3 shows the properties and methods of this class, and the online GitHub documentation provides example usage and benchmarks for accuracy, also shown in Fig. S7.

#### Helmholtz-Hodge decomposition of vector fields on dynamic surfaces

Our `DECLab` implementation includes a simple interface to generate a Helmholtz-Hodge decomposition of tangential vector fields [12], i.e. a decomposition into dilatational (curl-free), rotational (divergence-free), and harmonic parts. Let $v$ denote the 1-form field associated with the tangential surface velocity. In general, any surface velocity field can be decomposed in the following way

$$v = d\alpha + \delta\beta + h, \tag{13}$$

where $d$ is the exterior derivative, $\star$ is the Hodge star, and $\delta = \star d \star$ is the co-differential acting on 2-forms. The dilatational part of the velocity field, $d\alpha$, is given by the exterior derivative of the scalar potential $\alpha$. The rotational part of the velocity field, $\delta\beta$, is given by co-differential of the vector potential $\beta$ (confusingly $\beta$ is actually a 2-form despite the common naming convention). Finally, $h$ is a harmonic 1-form (i.e. $(d\delta + \delta d)h = 0$) encompassing the remaining aspects of $v$ that are neither dilatational nor rotational. Each term and potential function is given by the method `helmholtzHodgeDecomposition()`.

#### Lagrangian measures of time-integrated tissue strain

Endowing the evolving surface with a set of Lagrangian coordinates enables the construction of a *material metric*. The metric tensor, $\mathbf{g}(t)$, is a geometric object enabling the measurement of distances and angles between nearby points on the surface. The rate-of-deformation tensor describes how lengths and angles change locally as the surface deforms in time:

$$\frac{dg_{ij}(t)}{dt} = \nabla_i v_j + \nabla_j v_i - 2\, v_n\, b_{ij}, \tag{14}$$

where $v_i$, $i \in \{1, 2\}$, and $v_n$ denote the tangential and normal components, respectively, of the Lagrangian surface velocity, $\nabla_i$ denotes the covariant derivative with respect to the $i^{\text{th}}$ tangential coordinate, and $b_{ij}$ denote the components of the second fundamental form – another geometric tensor object that measures surface curvature. Essentially, Eq. (14) tells us that lengths and angles deform under the surface motion when there are gradients in the tangential velocity *and/or* when there is normal motion in curved regions of the tissue. We can then integrate the rate-of-deformation tensor along pathlines to construct a Lagrangian measurement of cumulative tissue strain, i.e.

$$\boldsymbol{\varepsilon}(t) = \frac{1}{2}\int_{\tau=0}^{\tau=t} d\tau\, \frac{d\mathbf{g}(\tau)}{d\tau} = \frac{1}{2}\left(\mathbf{g}(t) - \mathbf{g}(t_0)\right). \tag{15}$$

In the language of geometric elasticity, this is equivalent to the Green-St. Venant strain tensor [13], defined relative to the ‘undeformed’ reference configuration at time $t = 0$. The strain tensor can be decomposed into a dilatational (isotropic) component,

$$\frac{1}{2}\mathrm{Tr}\left[\mathbf{g}^{-1}(t_0)\boldsymbol{\varepsilon}(t)\right]\, \mathbf{g}(t_0), \tag{16}$$

and the deviatoric component

$$\mathrm{Dev}\left[\boldsymbol{\varepsilon}(t)\right] = \boldsymbol{\varepsilon}(t) - \mathrm{Tr}\left[\mathbf{g}^{-1}(t_0)\boldsymbol{\varepsilon}(t)\right]\mathbf{g}(t_0)/2. \tag{17}$$

### INTEGRATION OF TUBULAR WITH IMSANE

ImSAnE [14] is a tissue cartography package in MATLAB widely used in the developmental biology community. Since the aims of ImSAnE are a subset of those of TubULAR, we have implemented integration of the two toolkits in two ways: an ImSAnE Experiment class instance can be passed to TubULAR, or all of TubULAR’s functionality can be accessed entirely within ImSAnE. As shown in Fig. S8, we found existing methods within ImSAnE (and in other software) to be unable to automatically follow tissue-scale motions of a full tube-like surface.

#### Passing ImSAnE to TubULAR

An ImSAnE class instance can be passed to TubULAR as a first argument instead of a metadata struct. If no meshes have been found using ImSAnE already, then this step can be done with the newly created TubULAR instance, along with subsequent analysis. Alternatively, users may leverage any of ImSAnE’s surface detection methods before passing the result to TubULAR so long as the detection results in a series of meshes, named as indicated in the metadata passed to TubULAR. See the online documentation for further details.

#### Upgrades to ImSAnE integrate TubULAR’s functionality

ImSAnE workflows proceed in two parts: surface detection and surface fitting. ImSAnE now has an `integralDetector()` method for surface detection which mirrors TubULAR’s `getMeshes()` method. After surface detection, the `tubularFitter` creates a TubULAR instance and carries this object as a property. All functions can be accessed within the ImSAnE `Experiment` instance’s surfaceFitter.

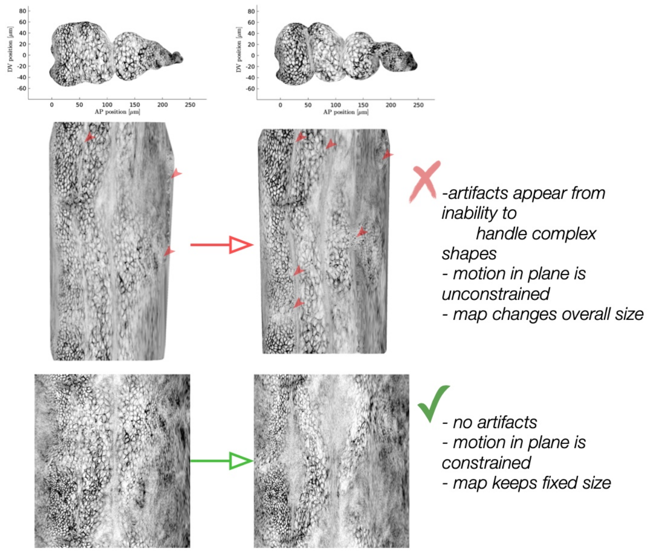


FIG. S8. **Previously existing methods fail to capture the full surface or follow tissue motion, as exemplified by a previously existing fitter in ImSAnE.** ImSAnE's `cylinderMeshWrapper` leads to image artifacts and self-intersections, as well as unconstrained motion of the tissue in the pullback plane, in contrast to our method. Red arrows indicate regions with parameterization problems in the pullback plane.

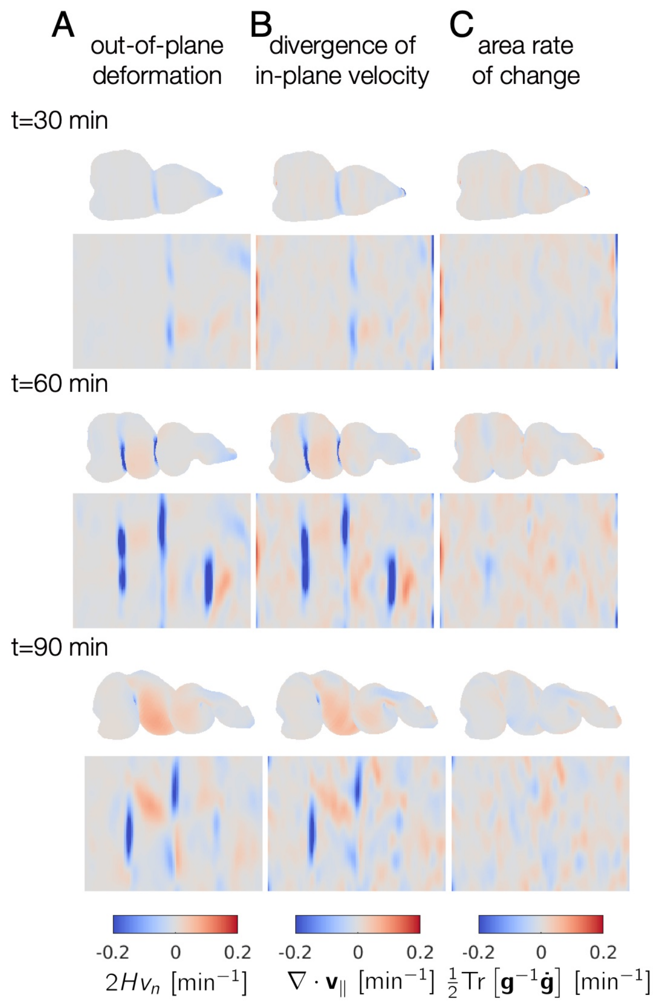


FIG. S9. **TubULAR measures the kinematic coupling between in-plane and out-of-plane motion, computing the rate of local area change across the organ, shown here for the developing midgut.** *(A)* The underlying out-of-plane deformation, defined as the normal motion $v_n$ times twice the mean curvature $H$, shows negative values at each constriction, where the mean curvature becomes negative. *(B)* DEC computation of the divergence of the in-plane velocity $\nabla \cdot \mathbf{v}_\parallel$ shows patterns of sinks in the constrictions and sources in the chambers' lobes, in synchrony with the out-of-plane deformation. *(C)* As a result of the match between in-plane and out-of-plane, the areal growth rate – defined as $\mathrm{Tr}\left[\mathbf{g}^{-1}\dot{\mathbf{g}}\right]/2$ – remains relatively quiescent.

### EXAMPLE OF INFERRING INTERCALATION RATES ('TISSUE TECTONICS') USING TUBULAR

Our approach aids in measurement of the contributions to tissue-scale convergent extension by cell shape change and oriented cell intercalation for dynamic curved surfaces. As illustrated in Fig. S10, in the absence of oriented cell divisions, cell shape change and oriented cell intercalations (or 'T1 transitions') both contribute to tissue-scale shear, in which different axes of the tissue converge and extend. An example of tracked cells demonstrating these two contributions is shown in Fig. S10. Note that each cell changes neighbors (through cell intercalations) and also changes its shape in a way to extend along the longitudinal (horizontal) axis and converge along the circumferential (vertical) axis.

As in [15], by imprinting the cell segmentation at time $t_0 = 0$ hr onto the surface and advecting the polygonal segmentation along material pathlines in 3D, we find a cell-by-cell measure of the tissue shear. This measures tissue shear by virtue of using a meso-scale measure of tissue flow from PIV in the pullback plane – providing a local measure of tissue-scale shear at each tissue patch on the surface. We can then directly compare the tissue-scale shear to the cell's actual shape change of cells segmented over time without explicitly tracking cells. The difference between the tissue-scale shear and the cell shape change is attributed to net oriented cell intercalations. Note that this method measures *net* intercalations, where one T1 event forming a new cell-cell junction aligned with the longitudinal direction can be canceled by a T1 event forming a new cell-cell junction aligned with the circumferential direction. Indeed, in the midgut many more cell intercalations occur than are measured by the net difference [15].

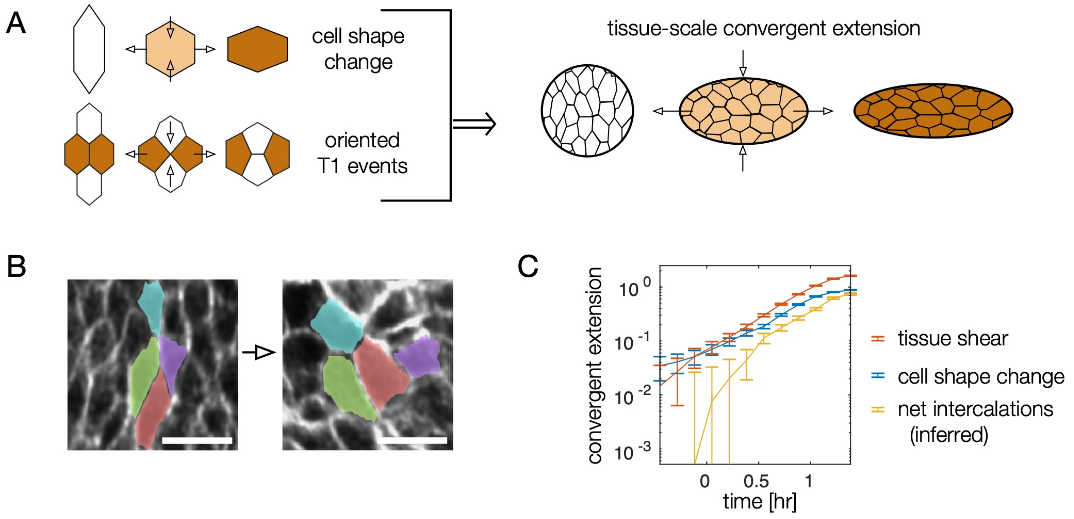


FIG. S10. **TubULAR aids in measurement of cell intercalations' contribution to tissue-scale convergent extension.** *(A)* In the absence of cell divisions, in-plane tissue-scale convergent extension occurs due to the changing shape of cells as well as the occurrence of oriented cell intercalations ('T1 events'). *(B)* During constrictions in the fly midgut, no cell divisions take place, but cells change shape and also intercalate in the endodermal layer. Scale bars are 10 $\mu$m. *(C)* We can then compare the cumulative effect of each contribution (blue and yellow) to the total tissue-scale convergent extension (orange, constricting along $\phi$ and extending along $s$). In the midgut endoderm, we directly measure tissue shear from the deviatoric component of the integrated strain computed from Lagrangian pathlines in 3D. This shear strain is almost entirely oriented along the longitudinal axis $s$. In order to compare directly this quantity to the cell shape change, we imprint the shapes of cells at $t = 0$ on the tissue surface, follow the outlines of these cells along tissue pathlines from coarse-grained particle image velocimetry measurements, and compute the cell shape anisotropy $(1 - a/b)\cos 2\theta$, where $a$ and $b$ are the semimajor and semiminor axes of the ellipse capturing each cell's moment of inertia tensor and $\theta$ is the cell's angle with respect to the material frame's longitudinal axis.

### ANALYSIS OF BEATING ZEBRAFISH HEART DYNAMICS

In the absence of cell proliferation, the relationship between local tissue area rate of change, in-plane divergence, and out-of-plane motion is [16]:

$$\nabla \cdot \mathbf{v}_{\|} - 2Hv_n = \frac{1}{2}\mathrm{Tr}\left[\mathbf{g}^{-1}\dot{\mathbf{g}}\right], \qquad (18)$$

where $\nabla \cdot \mathbf{v}_{\|}$ is the in-plane covariant divergence of the in-plane tissue velocities $\mathbf{v}_{\|}$, $H$ is the mean curvature of the surface, $v_n$ is the normal (out-of-plane) velocity, and $\mathrm{Tr}\left[\mathbf{g}^{-1}\dot{\mathbf{g}}\right]/2$ is the rate of local area change. We find the two terms on the left hand side are not equal, and in fact are anti-correlated. We measure their cross correlation between their circumferentially-averaged values, each of which is a function of the longitudinal coordinate, $s$, and of time, $t$: $\langle \nabla \cdot \mathbf{v}_{\|}\rangle_\phi(s,t)$ and $\langle 2Hv_n\rangle_\phi(s,t')$. Fixing the spatial coordinate but varying the time delay between measured values $\Delta = t - t'$ returns a sinusoidal correlation function parameterized by the time delay $\Delta$. This curve fits well to

$$C(\Delta) \approx A\cos\left(2\pi(\Delta - \tilde{\Delta})/T\right). \qquad (19)$$

The time shift corresponding to the maximum correlation – such that the in-plane and out-of-plane deformations would be in phase – is $\tilde{\Delta}$, which we report in the main text. This analysis gives insight into the kinematic properties of the tissue: tissue compressibility dominates the kinetics, prompting further modeling of the heart's mechanical cycle.

### A CYTOSKELETAL GEL ACTIVELY DEFORMS LIQUID DROPLETS

Figure 1L shows a snapshot of a DNA droplet in an active microtubule gel. Briefly, DNA droplets are assembled from multi-armed DNA nanostructures with self-interacting complementary overhangs [17]. Active flows are generated by microtubule filaments depleted through non-adsorbing polymers such as polyethylene glycol (PEG) and powered by clusters of kinesin motors. Kinesin motors convert chemical energy from the environment and generate inter-filament sliding [18]. The DNA droplets are covalently bound to kinesin motors resulting in mechanical coupling between the microtubule bundles and the surface of the droplets. Active stresses are exerted through the microtubule flows, generating droplet deformation and eventually eventually leading to pinch-off of the elongating neck and preventing coarsening of the DNA droplets. Further details of the experimental design and imaging are given in [19].

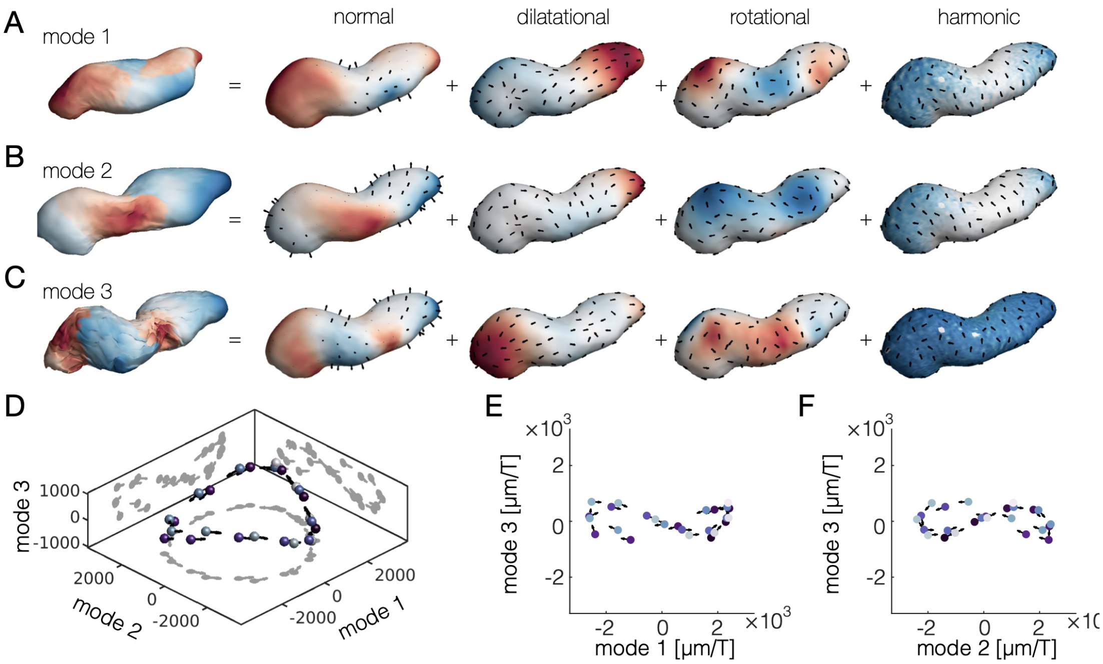


FIG. S11. **Higher order PCA modes contribute little to the description of the developing zebrafish heartbeat.** *(A-C)* Comparison of the first three PCA modes of heart beating dynamics shows that past the second mode, higher-order modes become noisy and do not capture directional pumping. *(D-F)* Projections of dynamics involving mode 3 (and higher modes) show nearly zero subtended area, unlike the dominant two modes, which sweep out a large area in mode space.

---

[1] E. Karzbrun, A. H. Khankhel, H. C. Megale, S. M. K. Glasauer, Y. Wyle, G. Britton, A. Warmflash, K. S. Kosik, E. D. Siggia, B. I. Shraiman, and S. J. Streichan, Nature **599**, 268 (2021).
[2] W. E. Lorensen and H. E. Cline, in *Proceedings of the 14th annual conference on Computer graphics and interactive techniques - SIGGRAPH '87*, Vol. 21 (ACM Press, New York, New York, USA, 1987) pp. 163–169.
[3] S. J. Osher and R. Fedkiw, *Level set methods and dynamic implicit surfaces.*, Applied mathematical sciences, Vol. 153 (Springer, 2003) pp. I–XIII, 1–273.
[4] P. Marquez-Neila, L. Baumela, and L. Alvarez, IEEE Transactions on Pattern Analysis and Machine Intelligence **36**, 2 (2014).
[5] T. F. Chan and L. A. Vese, IEEE Transactions on Image Processing **10**, 266 (2001).
[6] W. Zeng and X. D. Gu, *Ricci Flow for Shape Analysis and Surface Registration*, SpringerBriefs in Mathematics (Springer New York, New York, NY, 2013).
[7] N. Aigerman and Y. Lipman, ACM Transactions on Graphics **34**, 190:1 (2015).
[8] U. Pinkall and K. Polthier, Experimental Mathematics **2** (1993), 10.1080/10586458.1993.10504266.
[9] N. D. Cornea, D. Silver, and P. Min, IEEE Transactions on visualization and computer graphics **13**, 530 (2007).
[10] L. V. Ahlfors, *Lectures on Quasiconformal Mappings*, University lecture series (American Mathematical Society, 2006).
[11] T.-C. Lee, R. L. Kashyap, and C.-N. Chu, CVGIP: Graph. Models Image Process. **56**, 462–478 (1994).
[12] K. Crane, F. de Goes, M. Desbrun, and P. Schröder, in *ACM SIGGRAPH 2013 courses*, SIGGRAPH '13 (ACM, New York, NY, USA, 2013).
[13] E. Efrati, E. Sharon, and R. Kupferman, Journal of the Mechanics and Physics of Solids **57**, 762 (2009).
[14] I. Heemskerk and S. J. Streichan, Nature Methods **12**, 1139 (2015).
[15] N. P. Mitchell, D. J. Cislo, S. Shankar, Y. Lin, B. I. Shraiman, and S. J. Streichan, eLife **11**, e77355 (2022).
[16] M. Arroyo and A. Desimone, Physical Review E **79** (2009), 10.1103/physreve.79.031915.
[17] S. Biffi, R. Cerbino, F. Bomboi, E. M. Paraboschi, R. Asselta, F. Sciortino, and T. Bellini, Proceedings of the National Academy of Sciences **110**, 15633–15637 (2013).
[18] T. Sanchez, D. T. N. Chen, S. J. Decamp, M. Heymann, and Z. Dogic, Nature **491**, 431–434 (2012).
[19] A. M. Tayar, F. Caballaro, T. Anderberg, O. A. Saleh, M. C. Marchetti, and Z. Dogic, submitted (2022).
